# Direct and indirect effects: estimating the contributions along individual pathways during transient dynamics

**DOI:** 10.64898/2026.10.13.758872

**Authors:** Michael H. Cortez

## Abstract

All species in a community directly and indirectly affect each other, often through many causal pathways. The effects along each pathway can vary in magnitude as densities fluctuate during transient dynamics. Quantifying the changes in these pathway-specific effects is important for understanding the drivers of population-level dynamics. I present two computational methods that decomposes the total effect of one species on another into contributions along individual pathways during transient dynamics. These methods yield insight into which processes and species interactions are the primary drivers of transient population-level dynamics and how the effects of individual pathways scale with the number of intermediary species, time length, and interspecific interaction strength. I apply the method to two parameterized models, answering questions about the magnitudes of top-down effects in an intraguild predation system and the mechanisms driving disease reduction in a host-parasite system, and how the answers change through time.

## 1 Introduction

All species in an ecological community are directly or indirectly connected. The direct and indirect effects between species play a central role in shaping community structure and dynamics, for example, through trophic cascades (Pace et al., 1999; Shurin et al., 2002; Yang et al., 2010) and exploitative and apparent competition (Tilman, 1982; Holt and Lawton, 1994; Sanders et al., 2013; Holt and Bonsall, 2017). Moreover, because direct and indirect effects can be comparable in magnitude and operate on similar time scales (Dungan, 1986; Schoener, 1993; Menge, 1995, 1997; Montoya et al., 2009), understanding both is essential for explaining community dynamics.

The direct and indirect effects between two species can be viewed as distinct causal pathways through a community (Figure 1), where each pathway is defined by different sets of species or biological processes (Yodzis, 1988; Schoener, 1993; Wootton, 1994; Martins et al., 2024). Quantifying the contribution of each individual pathway is important because it identifies the specific processes or species interactions driving population-level dynamics (Connell, 1983*a*; Wootton, 1994, 2002). For example, the strongest pathways in food webs can determine which interspecific interactions are the most important for the propagation of trophic cascades (Wootton, 1994; Menge, 1995). Quantifying pathway-specific can also help determine which food-web interactions must be characterized to accurately predict the dynamics of a particular species (Yodzis, 1988; Menge, 1995). Importantly, the effects of each pathway can vary in sign and magnitude as species densities fluctuate, e.g., due to disturbance, species extinctions or invasions, and population oscillations. Thus, understanding transient population dynamics requires quantifying how pathway-specific effects vary through time.

**Figure 1:**
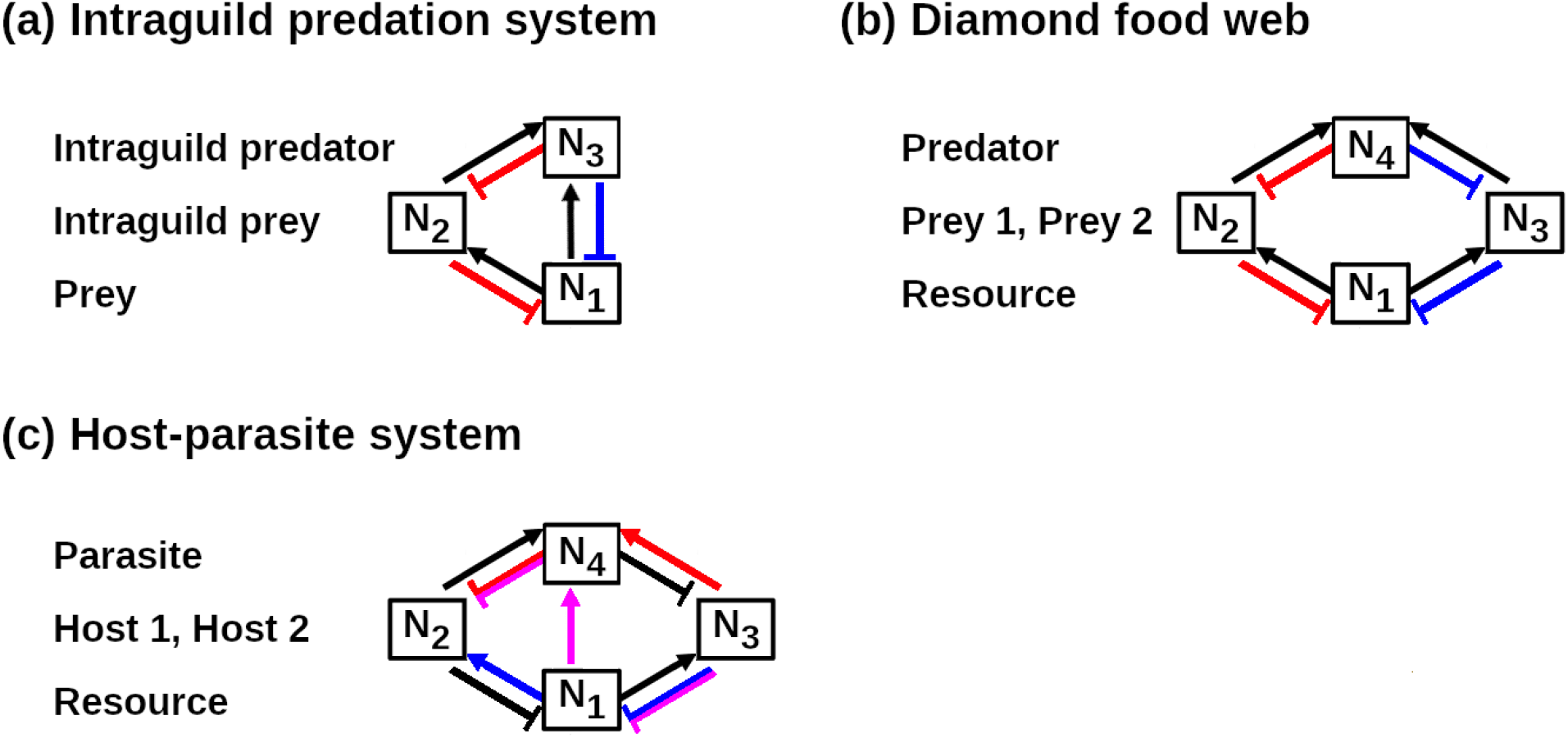
Visualizations of the multiple pathways connecting species in three simple interaction webs. (a) In the intraguild predation system, the intraguild predator affects the prey through a direct effect (blue) and an indirect effect (red) mediated by the intraguild prey. (b) In a diamond food web, the predator affects the resource through two indirect effects, one mediated by prey 1 (red) and one mediated by prey 2 (blue). (c) In the host-parasite system, host 2 affects host 1 through indirect effects mediated by the resource (blue), the parasite (red), and both the resource and parasite (magenta). In all panels, pointed and flat arrows indicate positive and negative direct effects between species, respectively.

Direct and indirect effects are empirically measured using species-removal experiments (Paine, 1966; Carpenter et al., 1987; Schmitz et al., 1997) and experimental press and pulse perturbations (Archibald, 1975; Bender et al., 1984; Menge, 1997; Carpenter et al., 2001). However, these approaches often measure the combined effect of many pathways.

For example, a resource’s response to manipulation of a top predator’s density will be due to the aggregate effect of the pathways involving each intermediate consumer (Figure 1b). In principle, the contributions of individual pathways can be estimated from carefully designed sets of perturbation experiments (Bender et al., 1984), but the number of experiments becomes impractical as the number of species and pathways increases. Thus, empirical studies have limited ability the quantity the contributions of individual pathways.

Parameterized process-based models and statistical models are alternative ways to estimate direct and indirect effects. Current computational and network-based methods can estimate the aggregate effect of all indirect pathways during transient dynamics, but not the effects of individual indirect pathways (Borrett et al., 2006, 2010; Burns et al., 2014). Approaches based on the Jacobian or community matrix compute pathway-specific effects, but the predictions from these methods are not guaranteed to apply to transient dynamics because the methods generally rely on systems being at or near equilibrium (Bender et al., 1984; Yodzis, 1988; Nakajima and Higashi, 1995; Yodzis, 1996; Takimoto et al., 2009; Abrams and Cortez, 2015). Statistical methods, like path analysis and structural equation models (Wootton, 1994; Gough and Grace, 1999; Grosholz et al., 2000), are also capable of estimating the effects of individual pathways. However, they become very data hungry when used to estimate temporally varying effects. In total, we currently lack a general theoretical framework for quantifying how the contributions of individual pathways change during transient community dynamics. That limits understanding of which processes drive observed dynamics and how that importance varies through time.

In this study, I present simulation-based methods for decomposing the total effect of one species on another into contributions arising from individual pathways during transient dynamics. I focus on density-mediated effects (but return to trait-mediated effects in the Discussion). Here, a direct effect is the physical impact of one species on the density of another (e.g., a predator reducing prey density) whereas an indirect effect is an impact mediated by changes in the densities of one or more intermediary species (e.g., a predator indirectly increasing resource density by reducing prey density). My approach extends a partitioning method from eco-evolutionary theory (Hairston et al., 2005; Ellner et al., 2011) and advances theory for density-mediated effects (Yodzis, 1988; Nakajima and Higashi, 1995; Burgess et al., 2013) by measuring how the signs and magnitudes of pathway-specific effects vary during transient dynamics. I apply the approach to parameterized models of an intraguild predation system and a host–parasite system, demonstrating how it can answer questions about which biological processes are driving observed population-level patterns and how the answers to those questions change through time.

## 2 Two methods for estimating individual effects during transient dynamics

My methods analyze time series from models of interacting species, with the goal of decomposing the total effect of one species on another into contributions from individual causal pathways.

The methods can be applied to models of structured and unstructured populations of any number of species (see “Method generality”). To facilitate understanding, I present the methods using a model of three unstructured populations. A general form for such a model is,

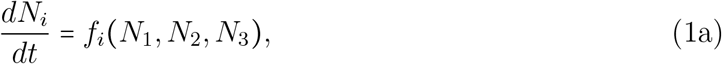

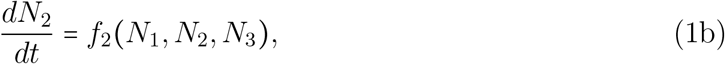

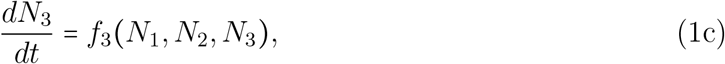

where *N*_*i*_ is the density and *f*_*i*_ *N*_1_, *N*_2_, *N*_3_ is the growth rate of species *i*.

To simplify the presentation, I focus on the effects of species 3 on species 1 (e.g., the top-down effects of an intraguild predator on a resource; Figure 1a). The total effect is the change in species 1’s density caused by a change in species 3’s density. This total effect is the sum of two pathways (Figure ??). Pathway 1 is a direct effect due to the growth rate of species 1 depending on species 3’s density (equation 1a). Pathway 2 is an indirect effect due to the growth rate of species 2 depending on species 3’s density (equation 1b) and the growth rate of species 1 depending on species 2’s density (equation 1a).

I present two methods for estimating the effects of each pathway from model simulations. Both methods start with the simulation of the original model where species 3 affects species 1 through both pathways. Then, using a factorial design, the methods compute the effect of each pathway using simulations from model variants corresponding to all combinations of the two pathways being present or absent. The two methods differ in how pathway absence is defined. Method 1 inactivates a pathway by holding species 3’s density fixed in model terms that are part of that pathway, thus preventing the effects of changes in species 3’s density from propagating along that pathway. This method is analogous to computing pathway-specific effects from short-term press perturbations that hold species 3’s density constant. Method 2 removes pathways by setting species 3’s density equal to zero in model terms that are part of that pathway. This method is analogous to computing pathway-specific effects from short-term removal experiments.

### 2.1 Method 1: Computing effects via pathway inactivation

Method 1 computes the effect of each pathway from simulations of four model variants corresponding to all combinations of the two pathways being active or inactive. Here, pathway inactivation means that species 3’s density is held fixed in model terms that are part of that pathway. The time series for the model variants are denoted,

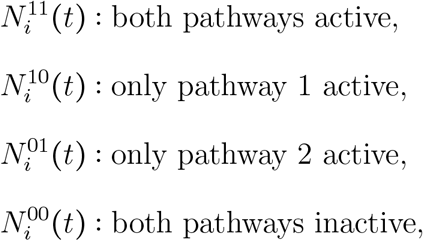

where the superscripts of 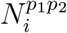 denote if pathway *j* is active (*p*_1_ =1, *p*_2_ =1) or inactive (*p*_1_ =0, *p*_2_ =0). Focusing on the time series for species 1, 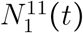 is identical to the observed time series from the original model where changes in species 3’s density affect species 1 through both pathways. 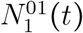 and 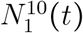 are counterfactuals where the changes in species 3’s density only affect species 1 through one pathway. 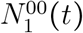 is the baseline for evaluating the contributions of each pathway because species 1’s dynamics are independent of changes in species 3’s density.

The steps for computing the pathway-specific effects from the model variants are the following. First, I simulate the original model over a set of time intervals, {(*t*_*s*−1_, *t*_*s*_)} (Figure 2a); the time series for the original model are denoted *N*_*i*_ (*t*).

**Figure 2:**
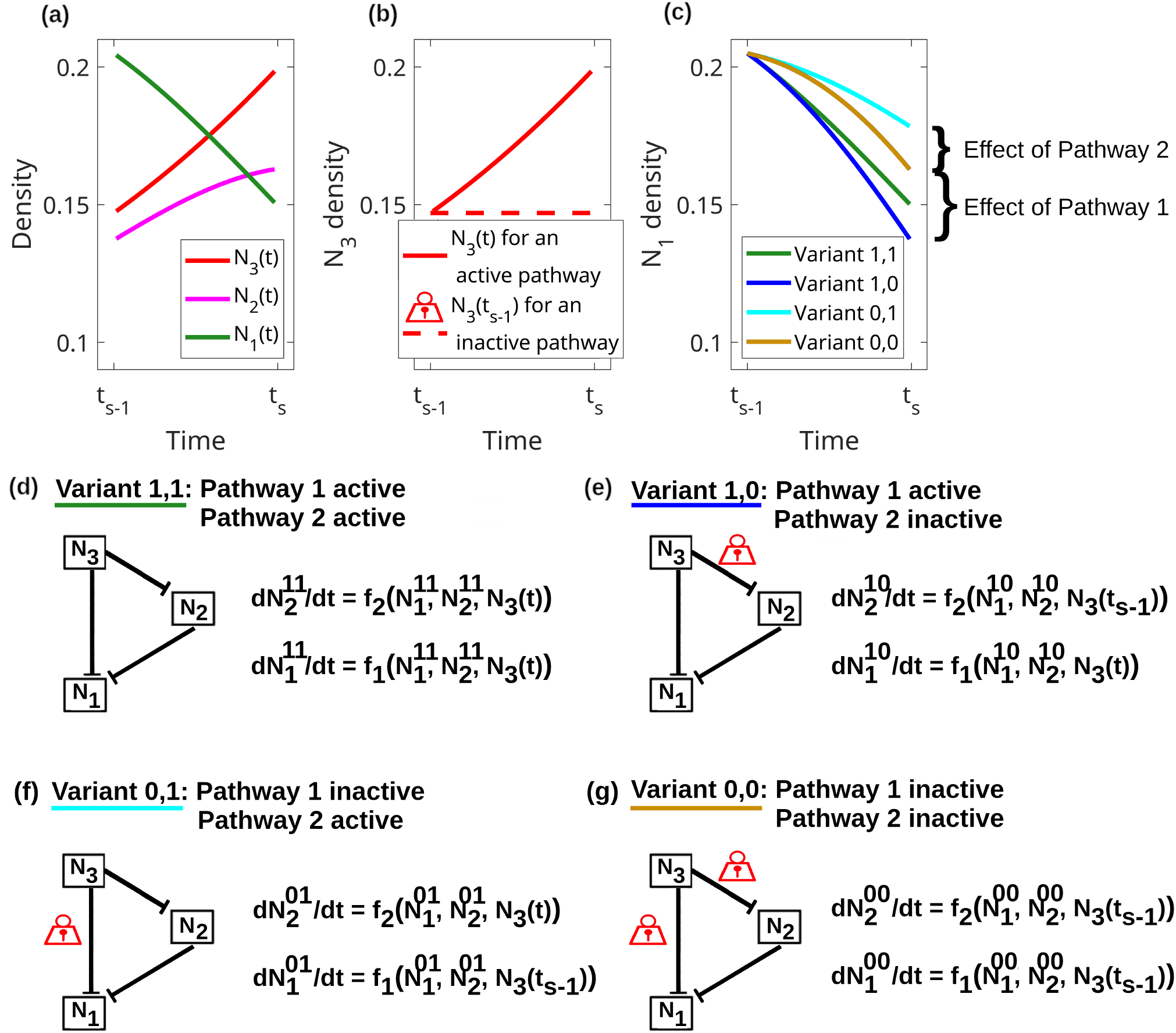
Steps used in Method 1 to compute the pathway-specific effects of species 3 on species 1 in a three-species model. (a) Time series for species 1 in green, species 2 in magenta, and species 3 in red, during the time interval (*t*_*s*−1_, *t*_*s*_). (b) In each model variant, the density of species 3 is identical to the original time series for active pathways (solid red) and held fixed (or “locked”) at its initial value for inactive pathways (dashed red). (c) Time series for species 1 from each model variant. The effect of each pathway is computed from the differences between the time series at the end of the interval. (d-g) Diagrams and equations for each model variant. In the diagrams, flat arrows denote negative direct effects between species and red locks denote the pathways that are inactivated by holding species 3’s density at its initial value. In each equation, the superscripts of 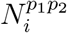 denote if pathway *j* is active (*p*_1_ = 1, *p*_2_ = 1) or inactive (*p*_1_ = 0, *p*_2_ = 0), where pathway 1 is the direct effect and pathway 2 is the indirect effect.

Second, each model variant is simulated over each interval in the following way.

1. The initial densities of all species are set equal to the observed state in the original model at time 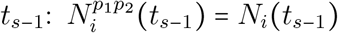
2. For each active pathway, species 3’s density is forced to be equal to the density in the original model in all model terms associated with that pathway: 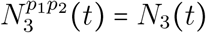 ; solid red curve in Figure 2b.
3. For each inactive pathway, species 3’s density is held fixed (or “locked’) at its initial density in all model terms associated with that pathway: 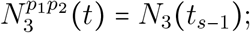 dashed red curve in Figure 2b.

Returning to the three-species model, the equations for the model variant with both pathways active (Figure 2d) are identical to those of the original model (1). Consequently, the time series for this model is identical to the original model (compare green curves in Figure 2ac). For the model variant where only pathway 1 is active (Figure 2e), pathway 1 is activated by forcing species 3’s density to be identical to the original model in species 1’s equation, 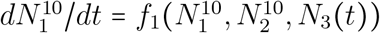. In contrast, pathway 2 is inactivated by holding species 3’s density fixed at its initial value in species 2’s equation, 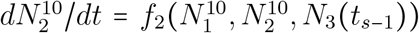 . Similarly, for the model variant where only pathway 2 is active (Figure 2e), pathway 2 is activated by forcing species 3’s density to be identical to the original model in species 2’s equation, 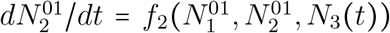 . Pathway 1 is inactivated by holding species 3’s density fixed at its initial value in species 1’s equation, 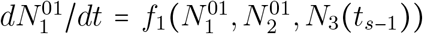 .For the model variant where both pathways are inactive (Figure 2f), species 3’s density is held fixed at its initial value in all terms of both equations, 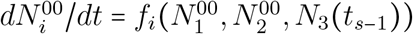 for *i* = 1, 2.

Finally, the effects of the pathways are computed from the differences between the model variants at the end of each interval (Figure 2c). Formally, the densities of the model variants are fit with the linear model,

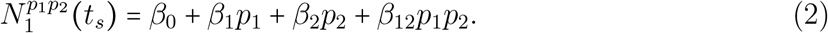

Here, *β*_*j*_ is the main effect of pathway *j, β*_12_ is the interactive effect of the pathways, and *p*_*j*_ acts as an indicator variable that has value 1 and 0 when pathway *j* is active and inactive, respectively. Solving the system of equations yields,

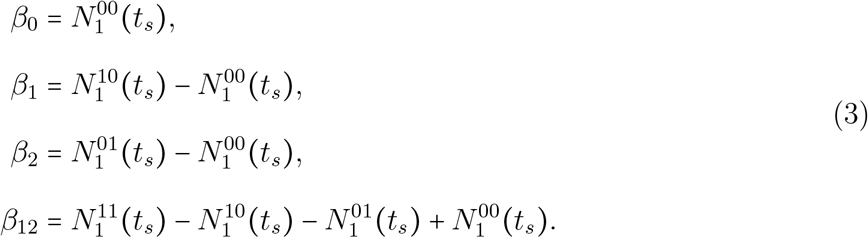

These formulas are identical to the main and interactive effects estimated by a two-way ANOVA. Dividing by the change in species 3’s density during the time interval yields the effect of each pathway, their interaction, and the total effect of species 3 on species 1,

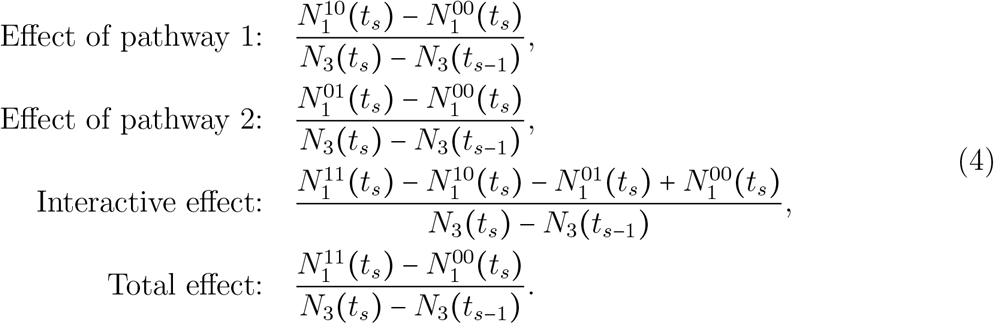

In these formulas, the individual effect of each pathway is the change in species 1’s density due to activation of just that particular pathway (braces in Figure 2c). The total effect is the change in species 1’s density due to activation of all pathways (difference between green and tan curves in Figure 2c).

In summary, Method 1 computes pathway-specific effects from model variants where the changes in species 3’s density only propagate along activated pathways. Applying the method to each time interval allows one to track how the pathway-specific effects change through time.

### 2.2 Factors affecting the effects measured with Method 1

What factors influence the magnitudes of the individual effects computed with Method 1? To answer this, I Taylor expand the general formulas for a direct effect and an indirect effect of species *n* on species 1 (Appendix S2). The leading order terms are

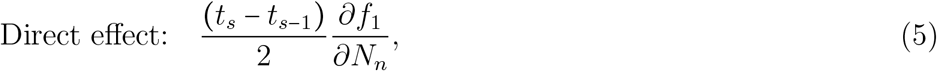

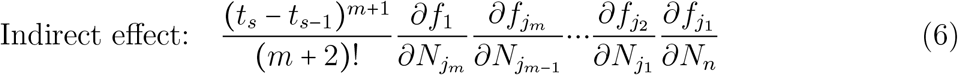

where species *j*_1_, …, *j*_*m*_ are the *m* intermediary species in the indirect pathway and all derivatives are evaluated at the initial densities for the time interval. These formulas show that the effect of each pathway depends on (i) interval length (*t*_*s*_−*t*_*s*−1_), (ii) pathway length (*m*), and (iii) the strengths of the interspecific interactions (*∂f*_*i*_/*∂N*_*j*_).

#### 2.2.1 Interval length

The choice of interval length, *t*_*s*_−*t*_*s*−1_, involves a trade-off between method accuracy and the ability to detect longer pathways. Formulas (4) are more accurate on shorter intervals. However, short intervals also reduce the effects of longer pathways involving many intermediary species. The intuitive reason is that indirect effects require more time to propagate along longer pathways because each species needs time to respond to changes in the density of the previous species in the pathway. Mathematically, the effect of a pathway containing *m* intermediary species is proportional to (*t*_*s*_−*t*_*s*−1_) ^*m*+1^, which increasingly downweights longer pathways as interval length decreases. Long intervals are also disadvantageous because they limit the detection of temporal variation in the pathways-specific effects. In addition, the interactive effects are larger in magnitude for longer intervals (Appendix S2), which complicates the interpretation of the individual effects.

To balance this trade-off, I measure the species’ growth rates (*f*_*i*_) on the time scale of the fastest changes in species densities and set the interval length to unity (*t*_*s*_−*t*_*s*−1_=1); see Appendices S1-S2 for details. With this scaling, an interval length of unity is short enough for the method to be accurate and long enough to avoid downweighting longer pathways.

#### 2.2.2 Pathway length

The effects of pathways decrease in magnitude as pathway length (*m*) increases. This is because the factor (*m*+2)! grows faster than exponentially and will ultimately dwarf the derivatives of the growth rates for sufficiently long pathways. All else being equal, this means pathways with many intermediary species have effects of smaller magnitude than pathways with few intermediary species.

#### 2.2.3 Interspecific interaction strengths

The effect of a pathway is larger when species in the pathway have stronger interspecific interactions. Direct effects are larger when species *n* has a stronger interspecific interaction with species 1 (larger *∂f*_1_/*∂N*_*n*_). Indirect effects are larger when consecutive species in the pathway have stronger interspecific interactions (e.g., larger *∂f*_*j2*_ /*∂N*_*j1*_).

This has two important implications. First, longer pathways need not have weaker effects than shorter pathways when some pathways contain especially strong or weak interspecific interactions. Second, the effects of pathways will vary through time because the strengths of the interspecific interaction depend on species densities. Consequently, the pathways and processes driving observed dynamics can differ as community composition changes.

#### 2.2.4 Similarity to prior equilibrium-based theory

Equations (5-6) are qualitatively similar to those from equilibrium-based approaches (Yodzis, 1988). The formulas from both approaches depend on the same products of derivatives, because those derivatives define the interspecific interaction strengths. However, there are two important differences. First, my formulas are evaluated at the different community states observed during the transient dynamics whereas equilibrium-based formulas are only evaluated at the model’s equilibrium. Second, my formulas for transient dynamics weight each pathway by the number of intermediary species and the interval length. In contrast, the equilibrium-based formulas weight the pathways by feedbacks between species at equilibrium because those indirect effects are measured over infinitely long time scales.

### 2.3 Method 2: Computing effects via pathway removal

Method 2 computes the effect of each pathway from simulations of four model variants corresponding to all combinations of the pathways being present or removed. Here, pathway removal means that species 3’s density is set equal to zero in all model terms in that pathway.

The steps for Method 2 are identical to Method 1 with two exceptions. First, for each removed pathway, species 3’s density is set to zero rather than held fixed at its initial density. Specifically, pathway 1 (the direct effect) is removed by setting species 3’s density to zero in the equation for species 1, e.g., 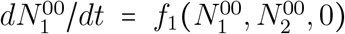 . Pathway 2 (the indirect effect) is removed by setting species 3’s density to zero in the equation for species 2, e.g., 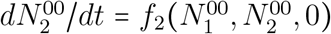 Second, the change in species 3’s density during a time interval is the difference between when it is present versus absent [*N*_3_ (*t*_*s*−1_) −0]; for simplicity, I use species 3’s density at the start of the interval to define its density when present.

The effects of each pathway, their interaction, and the total effect for Method 2 are

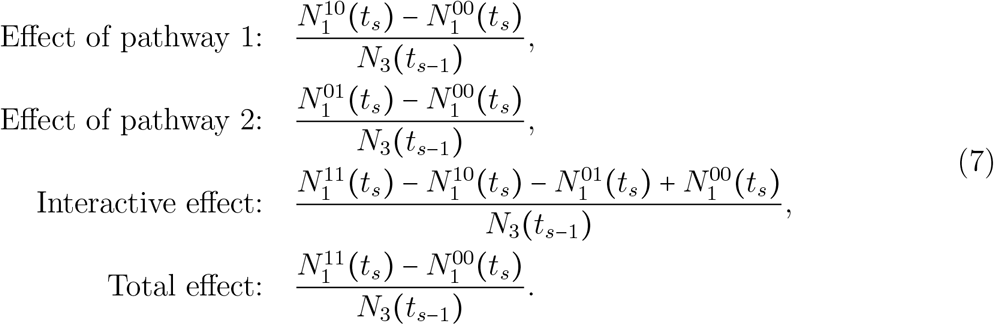

The individual effect of each pathway is the change in species 1’s density due to the presence of just that particular pathway. The total effect is the change in species 1’s density due to the presence of all pathways.

In summary, Method 2 computes pathway-specific effects using model variants where selected pathways are removed. Applying the method to each time interval allows one to track how the effect of each pathway changes through time.

### 2.4 Complementary benefits of using both methods

I recommend using both methods to compute the pathway-specific effects because they provide similar, but distinct, biological insight. Analogous to press perturbations, Method 1 provides insight about how the effects of small changes in species densities propagate through a community. This is useful for understanding how transient changes in the density of one species affect the transient dynamics of another species. In comparison, analogous to the removal experiment, Method 2 provides insight about how the presence of one species affects the transient dynamics of another species.

The two methods also have complementary advantages. The advantage of Method 1 is that the approximation formulas (5, 6) can help reveal why the pathway-specific effects vary through time (see the two applications to parameterized models). The advantage of Method 2 is that it avoids artificial spikes in effect sizes. Specifically, Method 1 has artificial spikes when the difference *N*_3_ (*t*_*s*_) − *N*_3_ (*t*_*s*−1_) in the denominators of formula (4) are small; this occurs whenever there is a peak or trough in the density of species 3. These artificial spikes are less likely with Method 2 because the divisors in formula (7) are the total density of species 3.

### 2.5 Method generality

Both methods extend in straightforward ways to models for more complex communities. First, the methods can be applied to models with any number of species. Second, for models with structured populations, the only minor conceptual difference is that each variable, *N*_*i*_, can represent the density of a population or subpopulation. Third, when the species are connected by *k*>2 pathways, the sole computational difference is that the methods use 2^*k*^ model variants (Appendix S3.1); these model variants correspond to the factorial combinations of the *k* pathways being present or absent. Forth, if multiple pathways overlap for the first few links (e.g., blue and magenta pathways in Figure 1c), applying the methods as described above will only compute the aggregate effect of the overlapping pathways. In Appendix S3.2, I show that the individual effects of each pathway can be computed by using model variants where the equations for particular species are duplicated. Altogether, these methods provide a general way for decomposing the total effect of one population on another into temporally varying contributions along any number of individual pathways.

## 3 Applications to parameterized models

I now apply both methods to parameterized models of an intraguild predation system and a host-parasite system. These examples illustrate how the methods can identify the main mechanisms through which species affect each other’s dynamics.

### 3.1 Example 1: Top-down effects in an intraguild predation system

Hiltunen et al. (2013) analyzed an intraguild predation model consisting of rotifers (the intraguild predator *Brachionus plicatilis*; *R*), flagellates (the intraguild prey *Oxyrrhis marina*; *F*), algae (the prey, *Chlorella autotrophica*; *A*), and a limiting substrate (nitrogen; *S*). The rotifer affects the algae through two pathways: a negative direct effect due to consumption of algae (Figure 1b) and a positive indirect effect due to the consumption of flagellates (Figure 1c). It is unknown how these two pathways combine to determine the total top-down effect of the predator during sustained oscillations. I use Method 1 and 2 to compute the strength of each pathway.

The model in Hiltunen et al. (2013) is

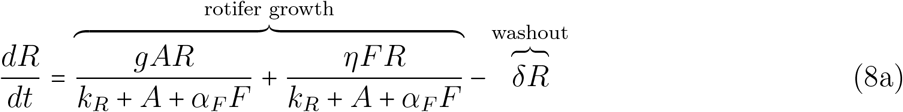

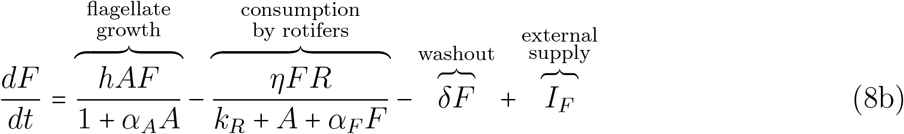

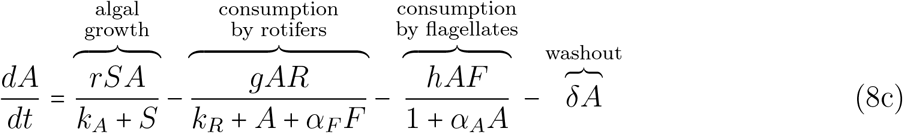

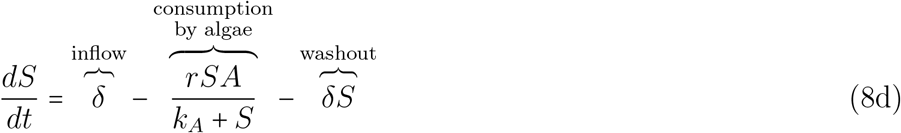

where the *k*_*i*_ are half-saturation constants, the *α*_*i*_ are handling-time parameters, *g, η, h* and *r* are maximum consumption rates, and *I*_*F*_ is the exogenous input rate of flagellates. The dilution rate, *δ*, determines the inflow rate for nutrients and the washout rate for all species. Additional details about model parameters, variants, and analyses are provided in Appendix S4.

Methods 1 and 2 show that the direct effect is larger than the indirect effect at all time points (10.5 and 5 times larger on average, respectively; Figure 3bd). My results suggest that this difference in magnitude is due to the indirect effect propagating along a longer pathway and flagellate densities typically being lower than algal densities. Using formulas (5-6), the ratio of the effect sizes is approximately,

**Figure 3:**
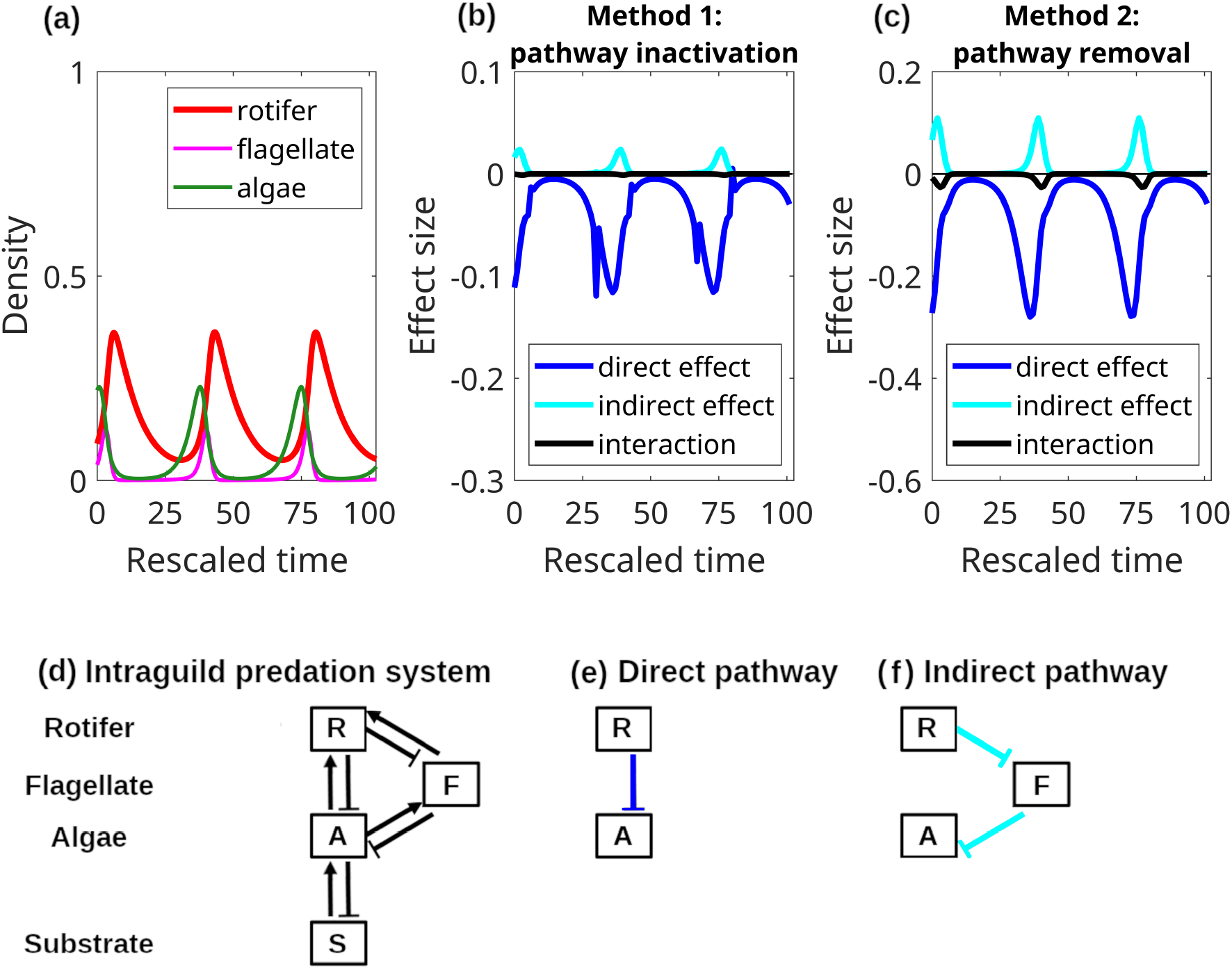
In an intraguild predation system, the negative direct effect of rotifers on algae is larger in magnitude than the positive indirect effect mediated by flagellates. (a) Time series for rotifer density (intraguild predator, *R*; red), flagellate density (intraguild prey, *F* ; magenta), and algal density (prey, *A*; green) generated by model (8); substrate density (*S*) dynamics are suppressed for clarity. Time series of the pathway-specific effects computed using (b) Method 1 and (c) Method 2. The curves are (blue) the direct effect, (cyan) the indirect effect mediated by the flagellate, and (black) the two-way interactive effect. The spikes in panel b are numerical artifacts due to small divisor issues at the peaks and troughs in rotifer density. Diagrams of the interspecific interactions in (d) the intraguild predation model, (e) the direct pathway between the rotifer and algae, and (f) the indirect pathway mediated by the flagellate. Pointed and flat arrows indicate positive and negative interactions, respectively.

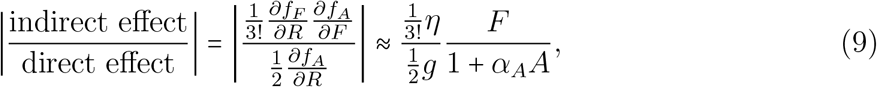

where *f*_*X*_ is the growth rate of species *X*; see Appendix S4.3 for details. This ratio is always less than 1 for two reasons. First, the rotifer has a larger direct effect on flagellates than algae because of a larger maximum consumption rate (*η* > *g*). However, that larger effect diminishes as it propagates along the longer indirect pathway (factor of 1/3! vs 1/2). This results in the first fraction being less than 1. Second, the direct and indirect effects are larger when algal density and flagellate density, respectively, are higher (e.g., note that the effect sizes peak when algal and flagellate densities are high). Because algal density is typically higher than flagellate density (Figure 3a), the second fraction is less than 1. Combined, this means the direct effect is larger than the indirect effect.

In total, Methods 1 and 2 predict that rotifers have a negative total effect on algae because the negative direct effect is greater than the positive indirect effect at all points in time.

### 3.2 Example 2: Mechanisms of disease reduction in a host-parasite system

Strauss et al. (2015) used a host-parasite model to study how the presence of a second host species (the water flea *Ceriodaphnia dubia*) affects the density of infected individuals in a primary host species (the water flea *Daphnia dentifera*). The parasite is a fungus (*Metschnikowia bicuspidata*) that is transmitted by spores released into the water. The total density of host 2 affects infected density in host 1 through five pathways: one mediated by spore uptake (Figure 1e), one mediated by spore release (Figure 1f), and three mediated by the shared algal resource (Figure 1g). The uptake-mediated pathway and two resource-mediated pathways have negative effects because they decrease infection rates by reducing spore density and the total density of host 1. The spore-release-mediated pathway and the third resource-mediated pathway have positive effects because they increase infection rates by raising spore density. While the observed total effect of the host 2 is negative (that is, a decrease in host 1 infected density), it is unknown which mechanisms(s) drive that outcome. Here, I use Methods 1 and 2 to compute the contributions along each pathway. However, to simplify the presentation, I compute the aggregate effect of the resource-mediated pathways; the independent effects are shown in Appendix S5.4.

The model in Strauss et al. (2015) includes class-structure because individual hosts can be susceptible or infected. To compute the effects of host 2 on host 1 infected density, I write the model in terms of the total density for each host (*N*_*i*_ for *i* =1, 2). The model equations are

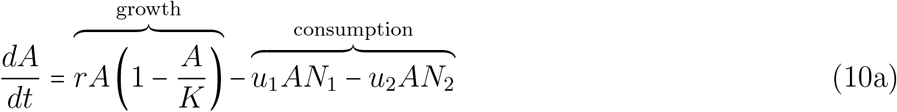

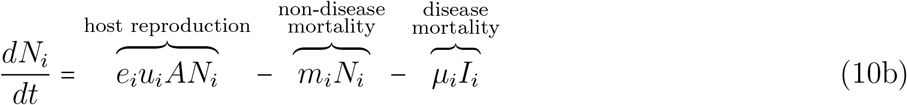

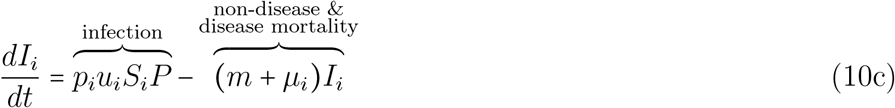

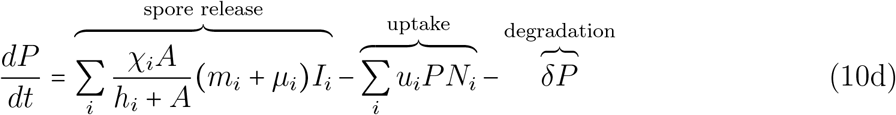

where *A* is algal density, *N*_*i*_, *I*_*i*_ and *S*_*i*_ *N*_*i*_ *I*_*i*_ are the total, infected, and susceptible densities for host *i* (*i* =1, 2), and *P* is spore density. Parameters *r* and *K* are the exponential growth rate and carrying capacity for the algae, *u*_*i*_ and *e*_*i*_ are host uptake (i.e., filtering) rates and conversion efficiencies, *m*_*i*_ and *µ*_*i*_ are non-disease and disease mortality rates, and *δ* is the spore degradation rate. Spore release occurs when infected individuals die, where *χ*_*i*_ is the maximum burst size (i.e., number of spores released per infected individual) and *h*_*i*_ is a resource-dependent half saturation constant. Additional details about model parameters, model variants, and analyses are provided in Appendix S5.

Methods 1 and 2 show that the uptake-mediated pathway has the largest effect on average, the aggregated resource-mediated pathways have a smaller effect, and the spore-release-mediated pathway has a much smaller effect (ratios of 2.03:1:0.07 for Method 1 and 1.47:1:0.07 for Method 2). Both methods also show that while initially smaller, the aggregated effect of the resource-mediated pathways is comparable in magnitude to the effect of the uptake-mediated pathway late in time (cyan and blue curves in Figure 4bc). This variation is explained by the dynamics of the susceptible and total densities of host 1. Using formula (6), the ratio of the effects is approximately,

**Figure 4:**
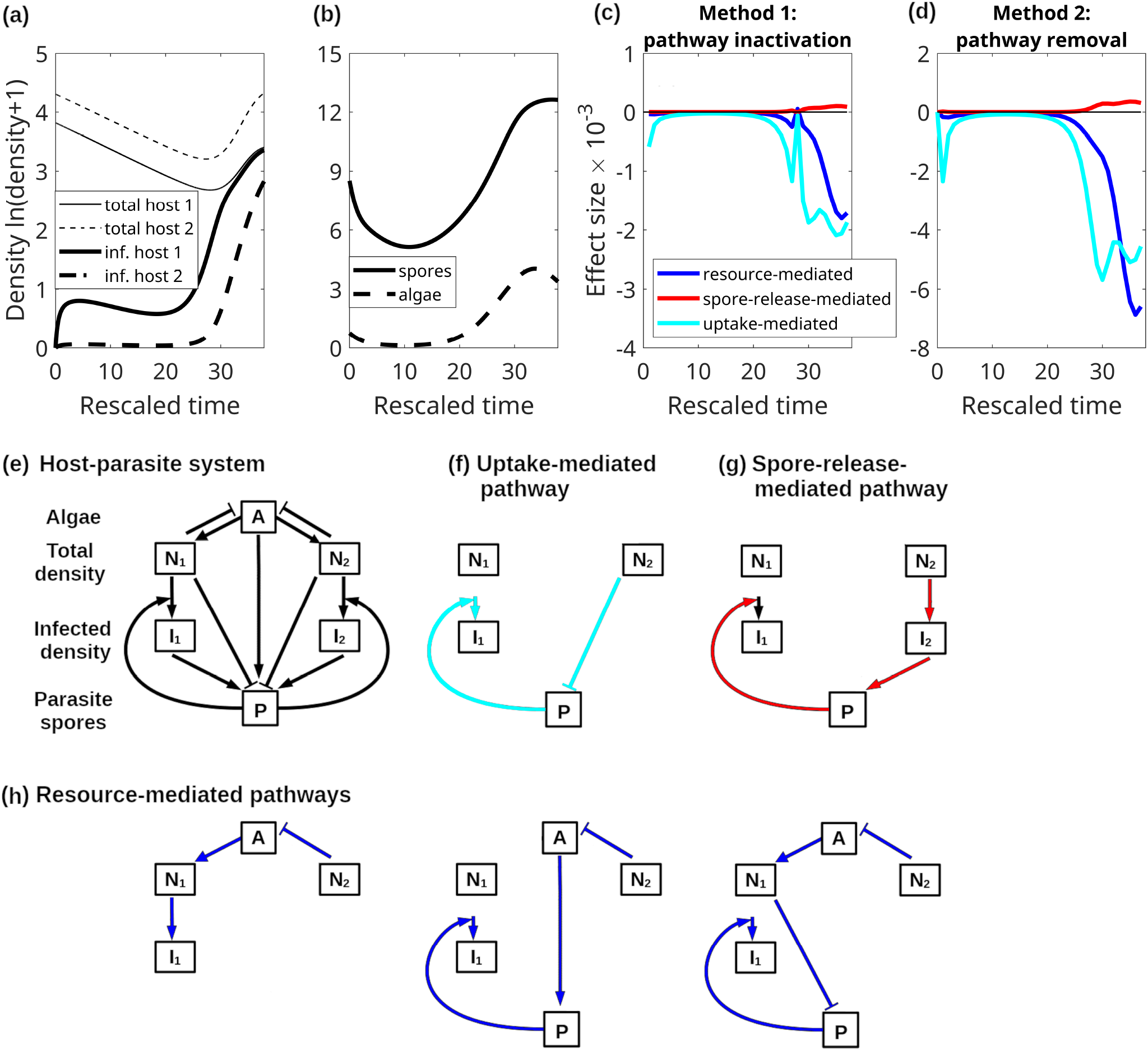
In the host-parasite system, the total effect of host 2 on host 1 infected density is initially dominated by the effect of the uptake-mediated pathway, but the uptake-mediated and the aggregate resource-mediated pathways have effects of comparable magnitude later in time. (a) Time series of total densities (thin curves) and infected densities (thick curves) for host 1 (solid curves) and host 2 (dashed curves) generated by model (10). (b) Time series of spore density (solid) and algal density (dashed). Time series of the effects computed using (c) Method 1 and (d) Method 2. The curves are (blue) the aggregate effect of the three resource-mediated pathways, (red) the effect of the spore-release-mediated pathway, and (cyan) the effect of the uptake-mediated pathway. The interactive effects are not shown because they are an order of magnitude smaller than the independent effects. The spikes in pale c are numerical artifacts due to small divisor issues at the troughs in host 2 density. Diagrams of the interspecific interactions in (e) the host-parasite model, (f) the uptake-mediated pathway, (g) the spore-release-mediated pathway, and (h) the three resource-mediated pathways. Pointed and flat arrows indicate positive and negative interactions, respectively.

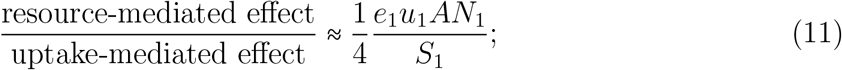

see Appendix S5.3 for mathematical details. The effect of the uptake-mediated pathway is proportional to susceptible host density, hence the *S*_1_ in the denominator. The aggregate effect of the resource mediated-pathways is proportional to host 1’s growth rate, hence the factor of *e*_1_*u*_1_*AN*_1_ in the numerator. The factor of 1/4 is due to the longer lengths of the resource-mediated pathways. Initially, the ratio is much less than 1 because algal density is low (*A* small; dashed curve in Figure 4b) and most host 1 individuals are susceptible (*S*_1_ = *N*_1_ − *I*_1_ ≈*N*_1_; difference between solid thick and solid thin curves in Figure 4a). However, the ratio is close to 1 at the end of the time series because host 1’s susceptible density is low and algal density is high. Consequently, the uptake-mediated pathway initially has a larger effect than the resource-mediated pathways, but the pathways have effects of comparable magnitude later in time.

In total, Methods 1 and 2 show that the negative effect of host 2 on host 1 infected density is primarily driven by spore uptake early in time and roughly equally driven by spore uptake and resource competition later in time. The positive effect of spore release by host 2 is minimal throughout.

## 4 Discussion

Direct and indirect effects propagate along many pathways through communities and are important for shaping the structure and dynamics of food webs (Paine, 1966; Pace et al., 1999; Yang et al., 2010), competitive systems (Connell, 1983*b*; Sanders et al., 2013; Holt and Bonsall, 2017), and mutualistic networks (Bascompte and Jordano, 2007; Okuyama and Holland, 2008). This study presented two methods that quantify which pathways have the strongest effects at different points in time. This is useful because it identifies the processes and species interactions driving observed dynamics, and how that changes through time. For instance, when applied to food web models, the methods can determine the fraction of a predator’s top-down effect that is due to each intermediate consumer and how those fractions change as species densities vary. Similarly, when applied to host-pathogen models, the methods can yield insight into which mechanisms drive changes in disease levels as host community composition changes (Keesing et al., 2006; Civitello et al., 2015; Cortez, 2024). For example, in the Strauss et al. (2015) model (10), disease reduction is initially driven by spore uptake, but competition for resources becomes equally important later in time (Figure 4). Consequently, the set of driving mechanisms changes even though the net effect has a constant sign. Thus, the analysis of individual pathways during transient dynamics can reveal not only which mechanisms drive patterns, but also how the relative importance of those mechanisms varies.

My methods generate testable predictions about the processes and species interactions that have the strongest effects on transient dynamics. Here I note four practical considerations when comparing theoretical results to empirical patterns. First, the methods require a model that is parameterized to the system and accurately reproduces it dynamics. This can require detailed knowledge of the system, including knowledge of the functional forms of species interactions (e.g., predator functional responses). As with any theoretical approach, prediction quality is constrained by how well the model represents the empirical system. Second, the computational costs of the methods grow rapidly with community complexity because the number of model variants scales exponentially with the number of pathways (2^*k*^). However, my analyses predict that the interactive effects of the pathways are smaller than their individual effects, suggesting that the individual effect can be accurately computed from the simulation of a much smaller number of model variants (see Appendix S3.1). Second, empirically testing predictions about the effects of individual pathways is likely to be difficult because pathways cannot be independently manipulated in most empirical systems. However, press perturbation or species-removal experiments could be used to test predictions about the aggregate effects of multiple pathways. Fourth, empirical sampling intervals are often longer than the unit interval lengths used in my methods. To facilitate theory-data comparisons, longer interval lengths can be used in both methods. However, this can complicate interpretation of the theoretical results because the magnitudes of the interactive effects increase with the interval length (Appendix S2).

A long-standing question in ecology is whether indirect effects attenuate along longer pathways, suggesting that only relatively short pathways influence ecological dynamics.

My methods can help address this by iteratively applying the method and estimating the individual direct and indirect effects of all species on a focal species. The approximation formula (6) shows how the magnitudes of the indirect effects scale with pathway length (an inverse factorial rate, 1/[*m*+2] !). This scaling is consistent with theoretical arguments that effects should attenuate as they propagate along longer pathways (Schoener, 1993; Wootton, 1994; Yodzis, 1996). At the same time, because the pathway-specific effects depend on interspecific interaction strengths, my results are also consistent with the argument that sufficiently strong interactions can cause long indirect pathways to remain important (Yodzis, 2000; Montoya et al., 2009). More generally, prior studies argue that short term dynamics are driven by short pathways and long term dynamics are driven by short and long pathways (Yodzis, 1996; Laska and Wootton, 1998). My methods provide a way to evaluate this argument by applying my transient-based methods to parameterized food web models and comparing predictions with equilibrium-based theories.

An important future application of my approach is the partitioning of trait-mediated effects along individual pathways. The trait-mediated effects of evolving and plastic traits can be as large as their density-mediated counterparts (Werner and Peacor, 2003; Preisser et al., 2005; Creel and Christianson, 2008; Hairston et al., 2005; Ezard et al., 2009). Current partitioning methods from eco-evolutionary dynamics estimate the total effect of species densities and species traits on ecological dynamics (Hairston et al., 2005; Ellner et al., 2011; Govaert et al., 2016; Govaert, 2018). Applying my pathway-centric approach to those methods would allow the quantification of both density-mediated and trait-mediated indirect effects along particular pathways. Such analyses would reveal which specific density and trait-based processes are driving eco-evolutionary dynamics.

In total, this study presents two methods for quantifying the pathway-specific effects between species during transient dynamics. It combines the strengths of equilibrium-based approaches for quantifying effects along particular pathways (Yodzis, 1988; Nakajima and Higashi, 1995) and computational approaches for quantifying aggregate affects during transient dynamics (Hairston et al., 2005; Ellner et al., 2011; Burns et al., 2014). By combining these strengths, the methods help answer questions about which mechanisms drive community dynamics and how the answers change through time.

## 5 Acknowledgments

I thank FSU for financial support, S.P. Ellner for helpful discussions, and M.L. Robertson, A.G. Dittmar, A. Chilton, and the members of FSU’s Ecology Reading Group for feedback on the paper.

