## Appendices for "Direct and indirect effects: estimating the contributions along individual pathways during transient dynamics"

Michael H. Cortez

### S1 Details about rescaling time

I rescale time in each model so that time is measured on the scale of the fastest changes in the species' densities. The steps for scaling the model are the following.

1. Compute the first and second derivatives of the density dynamics for each species, i.e., $dN_i/dt$ and $d^2N_i/dt^2$ for $i = 1, ..., n$.
2. Compute the ratio of the maximum values for each species, $R_i = \dfrac{\max_t\{d^2N_i/dt^2\}}{\max_t\{dN_i/dt\}}$.
3. Rescale time by the maximum ratio; the rescaled time variable is $\tau = R_{max}t$ where $R_{max} = \max_i R_i$.

This transformation scales time based on the time series with the steepest curvature relative to its largest slope. The reason for this scaling is that formula (4) approximations the total effect of species $n$ on species 1 using finite difference approximations of the species growth rates (i.e, their derivatives $dN_i/dt$). This rescaling bounds the error in the approximation such that the error is never greater than half of the maximum growth rate observed across all species. The following provides mathematical details. Note that to be increase the generality of the results, the following analysis of focuses on an $n$-species model and the total effect of species $n$ on species 1.

Generalizing equation (4) of the main text, the formula for the total effect of species $n$ on species 1 is

$$\text{total effect:}\quad \frac{N_1^{11}(t_s) - N_1^{00}(t_s)}{N_n(t_s) - N_n(t_{s-1})}. \tag{S1}$$

This formula relies on three finite difference approximations. To see this, recall that all model variants have the same initial conditions [i.e., $N_i^{p_1p_2}(t_{s-1}) = N_i(t_{s-1})$]. I write the formula for the total effect as

$$\frac{N_1(t_s) - N_1(t_{s-1})}{N_n(t_s) - N_n(t_{s-1})} = \frac{\dfrac{N_1^{11}(t_s) - N_1^{11}(t_{s-1})}{t_s - t_{s-1}} - \dfrac{N_1^{00}(t_s) - N_1^{00}(t_{s-1})}{t_s - t_{s-1}}}{\dfrac{N_n(t_s) - N_n(t_{s-1})}{t_s - t_{s-1}}} \tag{S2}$$

The first fraction in the numerator is a finite difference approximation of derivative of $N_1^{11}(t)$ during the time interval $(t_{s-1}, t_s)$. Specifically, approximating $N_1(t_s)$ using a Taylor expansion at $N_1(t_{s-1})$ yields

$$N_1^{11}(t_s) = N_1^{11}(t_{s-1}) + \left.\frac{dN_1^{11}}{dt}\right|_{t=t_{s-1}} (t_s - t_{s-1}) + \mathcal{O}([t_{s-1} - t_s]^2). \tag{S3}$$

Rearranging terms yields

$$\frac{N_1^{11}(t_s) - N_1^{11}(t_{s-1})}{t_s - t_{s-1}} = \left.\frac{dN_1^{11}}{dt}\right|_{t=t_{s-1}} + error. \tag{S4}$$

Thus, the first fraction in the numerator of equation (S2) is a finite difference approximation of the derivative $dN_1^{11}/dt$. Similarly, the other fractions in equation (S2) are finite difference approximations of $dN_1^{00}/dt$ and $dN_n/dt$,

$$\frac{N_1^{00}(t_s) - N_1^{00}(t_{s-1})}{t_s - t_{s-1}} = \frac{dN_1^{00}}{dt}\bigg|_{t=t_{s-1}} + error, \tag{S5}$$

$$\frac{N_n(t_s) - N_n(t_{s-1})}{t_s - t_{s-1}} = \frac{dN_n}{dt}\bigg|_{t=t_{s-1}} + error. \tag{S6}$$

The accuracies of the finite difference approximations depend on the second derivatives of the density dynamics. By Taylor's theorem, the magnitude of the higher order terms in equation (S2) is bounded above by $\frac{(t_s - t_{s-1})^2}{2}M$ where $M_1 = \max_{(t_{s-1},t_s)} |d^2N_1/dt^2|$ is the maximum magnitude of the second derivative of species 1's dynamics during the interval. This means that the error in equation (S3) is bounded above by

$$|error| \leq \frac{(t_s - t_{s-1})}{2}M_1. \tag{S7}$$

Similarly, the error in equation (S5) is bounded above by

$$|error| \leq \frac{(t_s - t_{s-1})}{2}M_n \tag{S8}$$

where $M_n = \max_{(t_{s-1},t_s)} |d^2N_n/dt^2|$ is the maximum magnitude of the second derivative of species $n$'s dynamics during the interval.

It would be ideal if time could be scaled such that the error terms are always smaller in magnitude than the derivatives at each point in time. However, this is impossible because the error will be non-zero at any point where the first derivative is zero and the curvature is non-zero. In particular, whenever species 1 or $n$ has a peak or trough in density, its derivative will be zero but its second derivative will be non-zero, implying that the error in the approximation is greater than the magnitude of the derivative.

My choice of rescaling ensures that if the interval size is unity, then the error terms are always less than half of the maximum rate of change observed for any species. To see this, let $\tau = R_{max}t$ be the dimensionless time variable, where $R_{max}$ is defined as in steps 1-3 above. Let $N_i(\tau)$ denote the time series for the dimensionless time variable. The finite difference approximation of the derivative of species $i$'s dynamics is

$$\frac{N_i(\tau_s) - N_i(\tau_{s-1})}{\tau_s - \tau_{s-1}} = \frac{dN_i}{d\tau}\bigg|_{\tau=\tau_{s-1}} + error \tag{S9}$$

where the error term satisfies $error \leq M(\tau_s - \tau_{s-1})/2$ for $M_i = \max_{(\tau_{s-1},\tau_s)} |d^2N_i/d\tau^2|$. Setting the interval size to unity ($\tau_s - \tau_{s-1} = 1$) and using the chain rule, the error in approximating species $i$'s derivative is bounded by

$$error \leq \frac{1}{2} \max_{(\tau_{s-1},\tau_s)} \left|\frac{d^2N_1}{d\tau^2}\right| = \frac{1}{2} \max_{(\tau_{s-1},\tau_s)} \left|\frac{d^2N_1}{dt^2}\left(\frac{dt}{d\tau}\right)^2\right| = \frac{1}{2R_{max}^2} \max_{(\tau_{s-1},\tau_s)} \left|\frac{d^2N_1}{dt^2}\right|. \tag{S10}$$

Also using the chain rule, the maximum growth rate observed across all species is

$$\max_{i,\tau}\left\{\frac{dN_i}{d\tau}\right\} = \max_{i,\tau}\left\{\frac{dN_i}{dt}\frac{dt}{d\tau}\right\} = \max_{i,t}\left\{\frac{dN_i}{dt}\right\}\frac{1}{R_{max}} \tag{S11}$$

Taking the ratio of the two previous equations and substituting the definition of $R_{max}$ yields

$$\frac{error}{\max_{i,\tau}\left\{\frac{dN_i}{d\tau}\right\}} \le \frac{1}{2R_{max}}\frac{\max_{(\tau_{s-1},\tau_s)}\left|\frac{d^2N_i}{dt^2}\right|}{\max_{i,t}\left\{\frac{dN_i}{dt}\right\}} \tag{S12}$$

$$= \frac{1}{2}\left[\frac{\max_{(\tau_{s-1},\tau_s)}\left|\frac{d^2N_i}{dt^2}\right|}{\max_{i,t}\left\{\frac{dN_i}{dt}\right\}}\right]\Bigg/\max_i\left\{\frac{\max_t\{d^2N_i/dt^2\}}{\max_t\{dN_i/dt\}}\right\} \le \frac{1}{2} \tag{S13}$$

Thus, if time is scaled as $\tau = R_{max}t$ and the interval size is chosen to be unity, then the error in the finite difference approximations are less than half of the maximum (rescaled) growth rate observed for any species.

### S2 Derivation of approximation formulas (5) and (6)

Here, I derive the formulas (5) and (6) from the main text. Those formulas assume an $n$-species model and are approximations for the pathway-specific effect of species $n$ on species 1. In the following, I first introduce some notation, define the Taylor series for the time series of each model variant, and then derive the approximation formulas. The end of the section presents approximation formulas for the interactive effects in special cases; the author was not able to derive a general formula that applied to any case.

**Notation:** Let $\vec{N}(t)$ denote the times series for the original model, $d\vec{N}/dt = \vec{f}(\vec{N})$. For given time interval $(t_{s-1}, t_s)$, let $\vec{N}_0 = \vec{N}(t_{s-1})$ denote the initial densities and $\vec{f}_0 = \vec{f}(\vec{N}_0) = [f_1(\vec{N}_0), f_2(\vec{N}_0), ..., f_n(\vec{N}_0)]$ denote the initial growth rates. Let $\mathbf{J}$ denote the Jacobian of the model evaluated at $\vec{N}_0$; its entries are $J_{ij} = \frac{\partial}{\partial N_j}\frac{dN_i}{dt}|_{\vec{N}_0} = \frac{\partial f_i}{\partial N_j}|_{\vec{N}_0}$. Let $H[\vec{f}_0, \vec{f}_0]$ denote the Hessian evaluated at $\vec{f}_0$; the entries of this vector are $H[\vec{f}_0, \vec{f}_0]_i = \sum_{j,k} \frac{\partial^2 f_i}{\partial N_j \partial N_k} f_j(\vec{N}_0) f_k(\vec{N}_0)$.

Let $N_i^{p_1p_2}(t)$ denote the time series for the four model variants, where $p_k = 1$ when pathway $k$ is active and $p_k = 0$ when pathway $k$ is inactive. It is important to recall three things. First, for each model variant, species $n$'s density is forced to be identical to the original model for active pathways, $N_n^{p_1p_2}(t) = N_n(t)$, whereas species $n$'s density is held at its initial value for the interval for inactive pathways, $N_n^{p_1p_2}(t) = N_n(t_{s-1})$. This means all model variants treat species $n$'s density as an exogenous variable and thus the dynamics of species 1 through $n-1$ do not influence the dynamics of species $n$. Second, all model variants have the same initial condition, i.e., $\vec{N}^{p_1p_2}(t_{s-1}) = \vec{N}(t_{s-1}) = \vec{N}_0$. Third, the initial growth rates for all model variants are the same, i.e., $d\vec{N}^{p_1p_2}/dt = \vec{f}_0$, because all models variants have identical initial conditions.

For each model variant, I denote the Jacobian and Hessian evaluated at $\mathbf{f}_0$ as $\mathbf{J}^{p_1p_2}$ and $H^{p_1p_2}[\vec{f}_0, \vec{f}_0]$. The Jacobian for each model variant, $\mathbf{J}^{p_1p_2}$, is similar to the Jacobian for the original model, $\mathbf{J}$, but there are two important differences. First, all entries in row $n$ are zero. This is because species $n$'s density is an exogenous variable. Second, each Jacobian has a zero in column $n$ for each inactivated pathway. All other entries of each Jacobian are identical to the Jacobian of the original model. The Hessian for each model variant, $H^{p_1p_2}[\vec{f}_0, \vec{f}_0]$, is also similar in structure to the Hessian for the original model, $H[\vec{f}_0, \vec{f}_0]$. Here, the key difference is that entry $i$ has zeros for every derivative with respect to $N_n$ if the corresponding pathway is inactivated.

To illustrate how the Jacobian and Hessian for each model variant differ from those for the original model, consider a three-species model where the model equations are $dN_i/dt = f_i(N_1, N_2, N_3)$ for $i = 1, 2, 3$, pathway 1 is the direct effect of species 3 on species 1, and pathway 2 is the indirect effect mediated by species 2. The Jacobian and Hessian for the original model and each model variant are,

$$\text{Original model} \qquad \mathbf{J} = \begin{bmatrix} J_{11} & J_{12} & J_{13} \\ J_{21} & J_{22} & J_{23} \\ J_{31} & J_{32} & J_{33} \end{bmatrix}, \quad H = \begin{bmatrix} \sum_{jk} \frac{\partial^2 f_1}{\partial N_j \partial N_k} f_j f_k \\ \sum_{jk} \frac{\partial^2 f_2}{\partial N_j \partial N_k} f_j f_k \\ \sum_{jk} \frac{\partial^2 f_3}{\partial N_j \partial N_k} f_j f_k \end{bmatrix}, \tag{S14}$$

Both pathways active

$$\mathbf{J}^{11} = \begin{bmatrix} J_{11} & J_{12} & J_{13} \\ J_{21} & J_{22} & J_{23} \\ 0 & 0 & 0 \end{bmatrix}, \quad H^{11} = \begin{bmatrix} \sum_{jk} \frac{\partial^2 f_1}{\partial N_j \partial N_k} f_j f_k \\ \sum_{jk} \frac{\partial^2 f_2}{\partial N_j \partial N_k} f_j f_k \\ 0 \end{bmatrix}, \tag{S15}$$

Only Pathway 1 active

$$\mathbf{J}^{10} = \begin{bmatrix} J_{11} & J_{12} & J_{13} \\ J_{21} & J_{22} & 0 \\ 0 & 0 & 0 \end{bmatrix}, \quad H^{10} = \begin{bmatrix} \sum_{jk} \frac{\partial^2 f_1}{\partial N_j \partial N_k} f_j f_k \\ \sum_{j,k\neq 3} \frac{\partial^2 f_2}{\partial N_j \partial N_k} f_j f_k \\ 0 \end{bmatrix}, \tag{S16}$$

Only Pathway 2 active

$$\mathbf{J}^{01} = \begin{bmatrix} J_{11} & J_{12} & 0 \\ J_{21} & J_{22} & J_{23} \\ 0 & 0 & 0 \end{bmatrix}, \quad H^{01} = \begin{bmatrix} \sum_{j,k\neq 3} \frac{\partial^2 f_1}{\partial N_j \partial N_k} f_j f_k \\ \sum_{jk} \frac{\partial^2 f_2}{\partial N_j \partial N_k} f_j f_k \\ 0 \end{bmatrix}, \tag{S17}$$

Both pathways inactive

$$\mathbf{J}^{00} = \begin{bmatrix} J_{11} & J_{12} & 0 \\ J_{21} & J_{22} & 0 \\ 0 & 0 & 0 \end{bmatrix}, \quad H^{00} = \begin{bmatrix} \sum_{j,k\neq 3} \frac{\partial^2 f_1}{\partial N_j \partial N_k} f_j f_k \\ \sum_{j,k\neq 3} \frac{\partial^2 f_2}{\partial N_j \partial N_k} f_j f_k \\ 0 \end{bmatrix}, \tag{S18}$$

where all quantities are evaluated at $\vec{N}_0$. For each Jacobian notice that (i) all entries in row 3 are zero because species 3 is treated as an exogenous variable, (ii) column 3 has zero entries in rows 1 and 2 whenever pathways 1 and 2, respectively, are inactivated because changes in species $n$'s density do not affect the dynamics of species 1 and 2, respectively, and (iii) all other entries are identical to those of $\mathbf{J}$ because the initial conditions of all model variants are identical to the initial conditions of the original model. For each Hessian notice that (i) entry 3 is always zero because species 3 is treated as an exogenous variable and (ii) the derivatives with respect to $N_n$ are absent in entries 1 and 2 when pathways 1 and 2 are inactive, respectively, because changes in species $n$'s density do not affect the dynamics of species 1 and 2, respectively.

**Taylor series:** With an abuse of notation, the Taylor series of the solutions for each model variant are,

$$\vec{N}^{11}(t_s) = \vec{N}_0 + \Delta \vec{f}_0 + \frac{\Delta^2}{2!}\mathbf{J}^{11}\vec{f}_0 + \frac{\Delta^3}{3!}\left([\mathbf{J}^{11}]^2\vec{f}_0 + H^{11}[\vec{f}_0, \vec{f}_0]\right) + \mathcal{O}(\Delta_4) \tag{S19}$$

$$\vec{N}^{10}(t_s) = \vec{N}_0 + \Delta \vec{f}_0 + \frac{\Delta^2}{2!}\mathbf{J}^{10}\vec{f}_0 + \frac{\Delta^3}{3!}\left([\mathbf{J}^{10}]^2\vec{f}_0 + H^{10}[\vec{f}_0, \vec{f}_0]\right) + \mathcal{O}(\Delta_4) \tag{S20}$$

$$\vec{N}^{01}(t_s) = \vec{N}_0 + \Delta \vec{f}_0 + \frac{\Delta^2}{2!}\mathbf{J}^{01}\vec{f}_0 + \frac{\Delta^3}{3!}\left([\mathbf{J}^{01}]^2\vec{f}_0 + H^{01}[\vec{f}_0, \vec{f}_0]\right) + \mathcal{O}(\Delta_4) \tag{S21}$$

$$\vec{N}^{00}(t_s) = \vec{N}_0 + \Delta \vec{f}_0 + \frac{\Delta^2}{2!}\mathbf{J}^{00}\vec{f}_0 + \frac{\Delta^3}{3!}\left([\mathbf{J}^{00}]^2\vec{f}_0 + H^{00}[\vec{f}_0, \vec{f}_0]\right) + \mathcal{O}(\Delta_4), \tag{S22}$$

where $\Delta = t_s - t_{s-1}$ is the interval length. The abuse of notation is that the formulas are only true for species 1 through $n-1$. For each formula, the equation for species $n$ should replaced with $N_n(t_s)^{p_1 p_2} = N_n(t_s)$ because species $n$ is treated as an exogenous variable. Note that the zero-order and first order terms are identical because the model variants have identical initial conditions and initial growth rates.

**Derivation of formula (5):** To derive formula (5), I assume pathway 1 is defined by a direct effect of species $n$ on species 1. I compute the difference,

$$N_1^{10}(t_s) - N_1^{00}(t_s) = \frac{\Delta^2}{2!} J_{1n} f_n + \frac{\Delta^3}{3!} \left( J_{11} J_{14} + \frac{\partial^2 f_1}{\partial N_n^2} f_n + 2 \sum_{i<n} \frac{\partial^2 f_1}{\partial N_i \partial N_n} f_i \right) f_n + \frac{1}{4!} \mathcal{O}(\Delta^4). \tag{S23}$$

where all quantities are evaluated at $\vec{N}_0$. I approximate the change in species $n$'s density with the finite difference $f_n = dN_n/dt = [N_n(t_s) - N_n(t_{s-1}]/[t_s - t_{s-1}]$, and divide by $[N_n(t_s) - N_n(t_{s-1})]$ to get

$$\frac{N_1^{10}(t_s) - N_1^{00}(t_{s-1})}{N_n(t_s) - N_n(t_{s-1})} = \frac{\Delta^2}{2!} J_{1n} + \frac{\Delta^3}{3!} \left( J_{11} J_{14} + \frac{\partial^2 f_1}{\partial N_n^2} f_n + 2 \sum_{i<n} \frac{\partial^2 f_1}{\partial N_i \partial N_n} f_i \right) + \frac{1}{4!} \mathcal{O}(\Delta^4). \tag{S24}$$

The lowest order term is the shortest pathway connecting species $n$ to species 1; substituting $J_{1n} = \frac{\partial f_1}{\partial N_n}$ yields formula (5) from the main text. The terms of the next highest order includes a pathway that revisits species 1 ($J_{11} J_{1n}$) and second order derivatives of species 1's growth rate.

**Derivation of formula (6):** To derive formula (6), I assume pathway 2 is defined by an indirect effect mediated one or more species. If the indirect effect is just mediated by species $j_1$, then I follow the same steps as above and compute,

$$\begin{aligned}\frac{N_1^{01}(t_s) - N_1^{00}(t_{s-1})}{N_n(t_s) - N_n(t_{s-1})} =& \frac{\Delta^2}{3!} J_{1j_1} J_{j_1 n} + \frac{\Delta^3}{4!} \Bigg( J_{11} J_{1j_1} J_{j_1 n} + J_{1j_1} J_{j_1 j_1} J_{j_1 n} \\ &+ \sum_{i \neq 1, j_1, n} J_{1i} J_{ij_1} J_{j_1 n} + \sum_{i \neq 1, j_1, n} J_{1j_1} J_{j_1 i} J_{in} \\ &+ J_{1j_1} \frac{\partial^2 f_{j_1}}{\partial N_n^2} f_n + 2 \sum_{i<n} J_{1j_1} \frac{\partial^2 f_{j_1}}{\partial N_i \partial N_n} f_i + 3 \sum_{i<n} J_{j_1 n} \frac{\partial^2 f_1}{\partial N_i \partial N_{j_1}} f_i \Bigg) + \mathcal{O}(\Delta^4),\end{aligned} \tag{S25}$$

where all quantities are evaluated at $\vec{N}_0$. The lowest order term is the shortest pathway connecting species $n$ to species 1 via species $j_1$. Terms of the next order include longer pathways that revisit a species (e.g., $J_{11} J_{1j_1}$ or $J_{1j_1} J_{j_1 j_1}$), longer pathways that include one other species (e.g., $J_{1j} J_{ij_1} J_{j_1 n}$ for $i \neq 1, j_1, n$), and second order derivatives of species 1's and species $j_1$'s growth rates. If the method is used to partition the effects along all possible pathways, then the higher order terms with additional species will be absent (i.e., the terms in the second line will not be present).

If the indirect effect is mediated by species $j_1$ and $j_2$, then the formula is,

$$\begin{aligned}\frac{N_1^{01}(t_s) - N_1^{00}(t_{s-1})}{N_n(t_s) - N_n(t_{s-1})} &= \frac{\Delta^3}{4!} J_{1j_2} J_{j_2 j_1} J_{j_1 n} \\ &+ \frac{\Delta^4}{5!} \left( \sum_i \left[ J_{1j} J_{ij_2} J_{j_2 j_1} J_{j_1 n} + J_{1j_2} J_{j_2 i} J_{ij_1} J_{j_1 n} + J_{1j_2} J_{j_2 j_1} J_{j_1 i} J_{in} \right] \right. \\ &\left. + J_{1j_2} J_{j_2 j_1} \frac{\partial^2 f_{j_1}}{\partial N_n^2} f_n + 2 \sum_{i<n} J_{1j_2} J_{j_2 j_1} \frac{\partial^2 f_{j_1}}{\partial N_i \partial N_n} f_i + 4 \sum_{i \neq j_1, n} J_{j_2 j_1} J_{j_1 n} \frac{\partial^2 f_1}{\partial N_i \partial N_{j_1}} f_i \right) + \mathcal{O}(\Delta^4).\end{aligned} \tag{S26}$$

Similar to the previous case, the lowest order term is the shortest pathway connecting species $n$ to species 1 via species $j_2$ and then species $j_1$. Terms of the next order include longer pathways that revisit a species, longer pathways that include one other species, and second order derivatives of species 1's and species $j_1$'s growth rates. As before, if the method is used to partition the effects along all possible pathways, then the higher order terms with additional species will be absent (i.e., the sum in the second line will only be over $i \in \{1, j_2, j_1\}$).

In general, for an indirect effect mediated by species $j_1, ..., j_m$, the leader order term is the ordered pathway connecting species $n$ to species 1, $\frac{\Delta^{m+1}}{(m+2)!} J_{1j_m} J_{j_m j_{m-1}} ... J_{j_2 j_1} J_{j_1 n}$ and the next higher order terms include longer pathways that revisit a species, longer pathways that include one other species, and second order derivatives of species 1's and species $j_1$'s growth rates. If the method is used to partition the effects along all possible pathways, then there will be no terms representing longer pathways that include one other species.

**Interactive effects:** I do not have a general formula for the interactive effects because the equations are too complex. In the following, I compute the 2-way interactive effects in a few specific cases. The formulas for those examples suggest that the 2-way interactive effects will typically be smaller in magnitude than the independent effects.

*Example 1:* Consider a 3-species model where pathway 1 is a direct effect ($N_3 \to N_1$) and pathway 2 is an indirect effect mediated one species ($N_3 \to N_2 \to N_1$); the Jacobian and Hessian for the original model and each model variant are shown in equations (S14)-(S18). For this model, the leading order terms for the individual and interactive effects are

$$\begin{aligned}\text{Pathway 1:} \quad & \frac{\Delta}{2!} J_{13}, \\ \text{Pathway 2:} \quad & \frac{\Delta^2}{3!} J_{12} J_{23}, \\ \text{Interaction:} \quad & \frac{3\Delta^3}{4!} J_{23} \frac{\partial^2 f_1}{\partial N_2 \partial N_3} f_3.\end{aligned} \tag{S27}$$

*Example 2:* Consider a 4-species models where pathway 1 is a direct effect ($N_4 \to N_1$) and pathway 2 is an indirect effect mediated by one species ($N_4 \to N_2 \to N_1$). The Jacobian for the full model is

$$J = \begin{pmatrix} J_{11} & J_{12} & J_{13} & J_{14} \\ J_{21} & J_{22} & J_{23} & 0 \\ J_{31} & J_{32} & J_{33} & J_{34} \\ J_{41} & J_{42} & J_{43} & J_{44} \end{pmatrix} \tag{S28}$$

For this model, the leading order terms for the individual and interactive effects are

$$\begin{aligned} \text{Pathway 1:} \quad & \frac{\Delta}{2!} J_{14}, \\ \text{Pathway 2:} \quad & \frac{\Delta^2}{3!} J_{13} J_{34}, \\ \text{Interaction:} \quad & \frac{3\Delta^3}{4!} J_{34} \frac{\partial^2 f_1}{\partial N_3 \partial N_4} f_4. \end{aligned} \tag{S29}$$

*Example 3:* Consider a 4-species models where pathway 1 is a direct effect ($N_4 \to N_1$) and pathway 2 is an indirect effect mediated by two species ($N_4 \to N_3 \to N_2 \to N_1$). The Jacobian for the full model is

$$J = \begin{pmatrix} J_{11} & J_{12} & 0 & J_{14} \\ J_{21} & J_{22} & J_{32} & 0 \\ J_{31} & J_{32} & J_{33} & J_{34} \\ J_{41} & J_{42} & J_{43} & J_{44} \end{pmatrix} \tag{S30}$$

For this model, the leading order terms for the individual and interactive effects are

$$\begin{aligned} \text{Pathway 1:} \quad & \frac{\Delta}{2!} J_{14}, \\ \text{Pathway 2:} \quad & \frac{\Delta^2}{4!} J_{12} J_{23} J_{34}, \\ \text{Interaction:} \quad & \frac{4\Delta^3}{5!} J_{23} J_{34} \frac{\partial^2 f_1}{\partial N_2 \partial N_4} f_4. \end{aligned} \tag{S31}$$

*Example 4:* Consider a 4-species models where pathways 1 and 2 are indirect effects mediated by one species ($N_4 \to N_2 \to N_1$ and $N_4 \to N_3 \to N_1$). The Jacobian for the full model is

$$J = \begin{pmatrix} J_{11} & J_{12} & 0 & J_{14} \\ J_{21} & J_{22} & J_{32} & 0 \\ J_{31} & J_{32} & J_{33} & J_{34} \\ J_{41} & J_{42} & J_{43} & J_{44} \end{pmatrix} \tag{S32}$$

For this model, the leading order terms for the individual and interactive effects are

$$\begin{aligned} \text{Pathway 1:} \quad & \frac{\Delta^2}{3!} J_{12} J_{24}, \\ \text{Pathway 2:} \quad & \frac{\Delta^2}{3!} J_{13} J_{34}, \\ \text{Interaction:} \quad & \frac{3\Delta^4}{5!} \left( J_{12} J_{34} \frac{\partial^2 f_2}{\partial N_4 \partial N_3} + J_{13} J_{34} \frac{\partial^2 f_3}{\partial N_4 \partial N_2} + 2 J_{24} J_{34} \frac{\partial^2 f_1}{\partial N_2 \partial N_3} \right) f_4. \end{aligned} \tag{S33}$$

Three lines of evidence suggest that the interactive effects will typically be smaller in magnitude than the independent effects. First, in all of the examples, the interactive effects have higher powers of $\Delta$ than the individual effects. All else being equal and assuming $\Delta = 1$, this suggests that the interactive effects are smaller in magnitude than the individual effects because the interactive effects are divided by larger factorials (e.g. $3/5!$ compared to $1/3!$ in example 4). Second, in some models, the mixed partial derivatives will be zero, implying that the leading order terms for the interactive effects are of even higher order. This occurs for models where the terms representing interspecific interactions only depend on the densities of the two species involved in the interspecific interaction. For example, all interspecific interaction terms in a Lotka-Volterra model, $dN_i/dt = r_i N_i + N_i \sum_j \alpha_{ij} N_j$, only depend on the densities of two species involved in the interaction. Third, the 2-way interactive effects are much smaller in magnitude than their corresponding independent effects for the two models analyzed in this study and other models analyzed by the author. In addition, the computed 3-way, 4-way, and higher interactive effects are even smaller in magnitude than the 2-way interactive effects in all of those models. Altogether, while it is unlikely that the interactive effects will always be smaller, the above suggests that it suggests that the interactive effects will typically be smaller than their corresponding individual effects.

### S3 Extensions of the methods

#### S3.1 Species with many connecting pathways

Methods 1 and 2 extend in a straightforward way to species with $k > 2$ connecting pathways. The only difference is that the methods require the simulation of $2^k$ model variants (but see the last paragraph of this subsection). Using a factorial design, these model variants correspond to all combinations of the $k$ pathways being present or absent (where pathway absence is either defined by inactivation or removal). The following provides the mathematical details. Note that to facilitate the presentation, the notation in this section differs from the notation of the main text.

I denote the time series for each model variant by $N_i^{\vec{v}}$ where the superscript $\vec{v}$ is the list of all present pathways. As examples, $N_i^1(t)$ is the time series from the model variant where only pathway 1 is present and all other pathways are absent; $N_i^{1,2}(t)$ is the time series from the model variant where only pathways 1 and 2 are present; and $N_i^{1,2,\ldots,k}(t)$ is the time series from the model variant where all pathways are present. I use $N_i^0(t)$ to denote time series from the model variant where all pathways are absent.

For each time interval, the final densities of the model variants are fit with the linear model,

$$N_1^{\vec{v}}(t_s) = \beta_0 + \sum_h \beta_h p_h + \sum_{h<i} \beta_{hi} p_h p_i + \sum_{h<i<j} \beta_{hij} p_h p_i p_j + \cdots . \tag{S34}$$

In this model, the $p_h$ are indicator variables that have values 1 and 0 when pathway $h$ is present or absent, respectively, The coefficient $\beta_h$ is the main effect of pathway $h$, $\beta_{hi}$ is the two-way interactive effect of pathways $h$ and $i$, $\beta_{hij}$ is the three-way interactive effect of pathways $h$, $i$, and $j$. The symbol $\cdots$ represents the sum of all other higher order interaction terms, e.g., the highest order term, $\beta_{1\ldots k} \prod_j p_j$, represents the $k$-way interaction of all pathways. Solving the system of equations for the coefficients yields

$$\begin{aligned}
\beta_0 &= N_1^0(t_s) \\
\beta_h &= N_1^h(t_s) - \beta_0 \\
\beta_{hi} &= N_1^{h,i}(t_s) - \beta_h - \beta_i - \beta_0 \\
\beta_{hij} &= N_1^{h,i,j}(t_s) - \beta_{hi} - \beta_{hj} - \beta_{ij} - \beta_h - \beta_i - \beta_j - \beta_0 \\
&\vdots
\end{aligned} \tag{S35}$$

The general pattern is that the coefficient for the independent effect of pathway $h$ is the difference between (a) the time series for the model variant where only pathway $h$ is present and (b) the time series for the model variant where all pathways are absent. The coefficient defining the interactive effect of pathways in list $\vec{v}$ is the difference between (a) the time series from the model variant where only the pathways in list $\vec{v}$ are present and (b) the sum of all lower order coefficients for subsets of $\vec{v}$.

Dividing the coefficients by the change in species $n$'s density yields the estimated individual effect of each pathway and the interactive effects of the pathways. To illustrate this, the following shows the algebraically simplified formulas for the total effect, individual effects, and two-way and three-way interactive effects.

$$\text{Total effect:} \quad \frac{\sum_{\vec{v}} \beta_{\vec{v}}}{N_n(t_s) - N_n(t_{s-1})} = \frac{N_1^{1,2,\ldots,k}(t_s) - N_1^0(t_s)}{N_n(t_s) - N_n(t_{s-1})}, \tag{S36}$$

$$\begin{array}{r}\text{Individual effect}\\ \text{of pathways } h\text{:}\end{array} \quad \frac{\beta_h}{N_n(t_s) - N_n(t_{s-1})} = \frac{N_1^h(t_s) - N_1^0(t_s)}{N_n(t_s) - N_n(t_{s-1})}, \tag{S37}$$

$$\begin{array}{r}\text{2-way interaction}\\ \text{for pathways } h, i\text{:}\end{array} \quad \frac{\beta_{hi}}{N_n(t_s) - N_n(t_{s-1})} = \frac{N_1^{hi}(t_s) - N_1^h(t_s) - N_1^i(t_s) + N_1^0(t_s)}{N_n(t_s) - N_n(t_{s-1})}, \tag{S38}$$

$$\begin{array}{r}\text{3-way interaction}\\ \text{for pathways } h, i, j\text{:}\end{array} \quad \frac{\beta_{hij}}{N_n(t_s) - N_n(t_{s-1})} = \frac{N_1^{hij} - N_1^{hi} - N_1^{hj} - N_1^{ij} + N_1^h + N_1^i + N_1^j - N_1^0}{N_n(t_s) - N_n(t_{s-1})}, \tag{S39}$$

where all values in the numerator of the last line are evaluated at $t_s$.

**Simulating model variants with many pathways:** The number of model variants ($2^k$) increases exponentially with the number of unique pathways ($k$) connecting the two species. This scaling is not costly for models with a small number of species, but it can be costly for models with many species. As noted in Appendix S2, I expect the 2-way and higher order interactive effects to be much smaller in magnitude than their corresponding independent effects; this is true for for all models presented in this study and others analyzed by the author. This suggests that a smaller number of model variants can be used to estimate the independent effects. The following presents two options.

The first option is to only simulate the one model variant where no pathways are present and the $k$ model variants where one pathway. In combination with the original model, these model variants are sufficient to compute the total effect and individual effects using equations (S36) and (S37). The disadvantage of this option is that this subset of model variants does not allow one to estimate the magnitudes of the interactive effects. However, one can estimate the sum of all interactive effects by (i) computing the sum of all individual effects by summing equation (S37) across all pathways, (ii) computing the total effect using equation (S36), and (iii) computing the difference between those two quantities. If the difference is smaller in magnitude than any individual effect, then this suggests that the interactive effects are likely smaller than the individual effects. Alternatively, if the difference is larger in magnitude than an individual effect, then this suggests that the interactive effects may not be negligible and additional model variants should be simulated to compute those interactive effects.

The second option is to simulate the one model variant where no pathways are present, the $k$ model variants where only one pathway is present, and the $k(k-1)$ pathways where two pathways are present. This set of $k^2 + 1$ model variants is sufficient to compute the total effect, individual effects, and 2-way interactive effects using equations (S36)-(S38). While this option requires more model variants, it has the advantage that one can directly determine if any 2-way interactive effects are comparable in magnitude to their corresponding individual effects. In addition, similar to before, one can estimate the sum of all higher order interactive effects by (i) computing the sum of all individual and two-way effects by summing equations (S37)-(S38) across all pathways, (ii) computing the total effect using equation (S36), and (iii) computing the difference between those two quantities. If difference is smaller in magnitude than any individual effect, then this suggests that the higher order interactive effects are likely small in magnitude. If instead the difference is larger in magnitude, then this suggests that the higher order interactive effects may not be negligible and additional model variants should be simulated to compute those interactive effects.

#### S3.2 Species with overlapping pathways

Methods 1 and 2 can be applied to species with pathways that initially overlap. For such cases, the key differences is that particular model equations are duplicated so that the overlapping pathways can be separated. The methods are then applied to the extended system as before.

The general steps for creating the extended system of equation are the following. Assume a pair of species has $k$ pathways whose first $m$ links are the same.

1. For each of the $m$ overlapping links, $N_d \to N_r$, let $N_d$ denote the donor species and $N_r$ denote the recipient species. Let $N_{r,j}$ denote the $j$th copy of the density for the recipient species ($j = 1, ..., k$).

2. Create $k$ copies of the equation for each recipient species. In the $j$th copy of the equation, replace all instances of $N_r$ with $N_{r,j}$.

3. In the equations for the other species, replace $N_r$ with $N_{r,j}$ in each model term that is in the $j$th pathway.

In the following, I illustrate these steps and the construction of the model variants using 2 examples.

**Example 1:** Consider the intraguild predation system in Figure 1a and focus on the bottom-up effect of the substrate ($S$) on the rotifer ($R$). The equations for the original model have the form,

$$
\begin{aligned}
\text{Rotifer} \quad \frac{dR}{dt} &= f_R(R, F, A), \\
\text{Flagellate} \quad \frac{dF}{dt} &= f_F(R, F, A), \\
\text{Algae} \quad \frac{dA}{dt} &= f_A(R, F, A, S), \\
\text{Substrate} \quad \frac{dS}{dt} &= f_S(A, S).
\end{aligned}
\tag{S40}
$$

The bottom-up effect is composed of two indirect pathways: pathway 1 is mediated by just the algae ($S \to A \to R$) and pathway 2 mediated by the algae and flagellate ($S \to A \to F \to R$). Importantly, both pathways share the initial link from the substrate to the algae ($S \to A$). Applying Methods 1 and 2 as described in the main will not estimate the individual effects of the two pathways because holding the substrate density fixed (at its initial value or zero) will inactivate/remove both pathways. Thus, I create an extended system by following steps 1-3 from above.

*Step 1:* The two pathways share an initial link from the substrate (donor species) to the algae (recipient species). Let $A_1$ and $A_2$ denote the two copies of the algal density.

*Steps 2 & 3:* The extended system of equations with two copies of the algal equation is,

$$\begin{aligned}
\text{Rotifer} \quad & \frac{dR}{dt} = f_R(R, F, A_1), \\
\text{Flagellate} \quad & \frac{dF}{dt} = f_F(R, F, A_2), \\
\text{Algae-1} \quad & \frac{dA_1}{dt} = f_A(R, F, A_1, S), \\
\text{Algae-2} \quad & \frac{dA_2}{dt} = f_A(R, F, A_2, S), \\
\text{Substrate} \quad & \frac{dS}{dt} = f_S(A, S).
\end{aligned} \tag{S41}$$

I note two things about this extended system of equations. First, if all pathways are present, then the dynamics of $A_1$ and $A_2$ are identical to the dynamics of $A$ in the original model (S40). Second, the rotifer growth rate only depends on $A_1$ whereas the flagellate growth rate only depends on $A_2$. This matches how algal density affects the model terms along each pathway. It also separates the two pathways: pathway 1 is mediated by algae-1 ($S \to A_1 \to R$) whereas pathway 2 is mediated by algae-2 and the flagellate ($S \to A_2 \to F \to R$).

The individual effects of the two pathways are estimated by applying Methods 1 and 2 to variants of the extended system. For example, if applying Method 2, then the equations for the model variants are,

$$\begin{array}{lll}
\text{Pathway 1 present} & \text{Pathway 1 removed} & \text{Both removed} \\
\text{Pathway 2 removed} & \text{Pathway 2 present} & \\
\frac{dR^{10}}{dt} = f_R(R^{10}, F^{10}, A_1^{10}), & \frac{dR^{01}}{dt} = f_R(R^{01}, F^{01}, A_1^{01}), & \frac{dR^{00}}{dt} = f_R(R^{00}, F^{00}, A_1^{00}) \\
\frac{dF^{10}}{dt} = f_F(R^{10}, F^{10}, A_2^{10}), & \frac{dF^{01}}{dt} = f_F(R^{01}, F^{01}, A_2^{01}), & \frac{dF^{00}}{dt} = f_F(R^{00}, F^{00}, A_2^{00}) \\
\frac{dA_1^{10}}{dt} = f_A(R^{10}, F^{10}, A_1^{10}, S(t)), & \frac{dA_1^{01}}{dt} = f_A(R^{01}, F^{01}, A_1^{01}, 0), & \frac{dA_1^{00}}{dt} = f_A(R^{00}, F^{00}, A_1^{00}, 0), \\
\frac{dA_2^{10}}{dt} = f_A(R^{10}, F^{10}, A_2^{10}, 0), & \frac{dA_2^{01}}{dt} = f_A(R^{01}, F^{01}, A_2^{01}, S(t)), & \frac{dA_2^{00}}{dt} = f_A(R^{00}, F^{00}, A_2^{00}, 0),
\end{array}$$

where $S(t)$ is treated as an exogenous variable whose values are identical to those in the original model. The variant with both pathways removed (right column) is the base case where the substrate has no effect on the rotifer. The other two variants (left and middle columns) represent counterfactuals where the substrate indirectly affects the rotifer only through one pathway. The equations for Method 1 are similar, with the only difference being that the zeros are replaced with the initial substrate density for the time interval, i.e., $S(t_{s-1})$.

**Example 2:** To illustrate the steps when the pathways have multiple overlapping links, assume the substrate is a biotic organism that feeds on another species with density $Q$. The equations for this new model are

$$\begin{aligned}
\text{Rotifer} \quad & \frac{dR}{dt} = f_R(R, F, A), \\
\text{Flagellate} \quad & \frac{dF}{dt} = f_F(R, F, A), \\
\text{Algae} \quad & \frac{dA}{dt} = f_A(R, F, A, S), \\
\text{Substrate} \quad & \frac{dS}{dt} = f_S(A, S, Q), \\
\text{Q-species} \quad & \frac{dQ}{dt} = f_Q(S, Q).
\end{aligned} \tag{S42}$$

For this model, the bottom-up effect of species $Q$ on the rotifer is composed of two pathways: pathway 1 is mediated by the substrate and algae ($Q \to S \to A \to R$) and pathway 2 mediated by the substrate, algae, and flagellate ($Q \to S \to A \to F \to R$). Both pathways share two links ($Q \to S$ and $S \to A$). Thus, I create an extended system by following steps 1-3.

*Step 1:* We must duplicate the variables for both the substrate (because it is the recipient in the link $Q \to S$) and the algae (because it is the recipient in the link $S \to A$). Let $S_1$, $S_2$, $A_1$, and $A_2$ denote the two copies of the substrate and algal density.

*Steps 2 & 3:* The extended system of equations is,

$$\begin{aligned}
\text{Rotifer} \quad & \frac{dR}{dt} = f_R(R, F, A_1), \\
\text{Flagellate} \quad & \frac{dF}{dt} = f_F(R, F, A_2), \\
\text{Algae-1} \quad & \frac{dA_1}{dt} = f_A(R, F, A_1, S_1), \\
\text{Algae-2} \quad & \frac{dA_2}{dt} = f_A(R, F, A_2, S_2), \\
\text{Substrate-1} \quad & \frac{dS_1}{dt} = f_S(A_1, S_1, Q), \\
\text{Substrate-2} \quad & \frac{dS_2}{dt} = f_S(A_2, S_2, Q),
\end{aligned} \tag{S43}$$

First, if all pathways are present, then the dynamics of $S_1$ and $S_2$ are identical to the dynamics of $S$ in the original model and $A_1$ and $A_2$ have dynamics identical to those of $A$ in the original model (S42). Second, the duplication of the substrate and algal equations separates the pathways: pathway 1 is mediated by substrate-1 and algae-1 ($Q_1 \to S_1 \to A_1 \to R$) and pathway 2 is mediated by substrate-2, algae-2, and the flagellate ($Q_2 \to S_2 \to A_2 \to F \to R$).

The individual effects of the two pathways are estimated by applying Methods 1 and 2 to variants of the extended system. For example, if applying Method 2, then the equations for the model variants are,

$$
\begin{array}{lll}
\text{Pathway 1 present} & \text{Pathway 1 removed} & \text{Both removed} \\
\text{Pathway 2 removed} & \text{Pathway 2 present} & \\
\frac{dR^{10}}{dt} = f_R(R^{10}, F^{10}, A_1^{10}), & \frac{dR^{01}}{dt} = f_R(R^{01}, F^{01}, A_1^{01}), & \frac{dR^{00}}{dt} = f_R(R^{00}, F^{00}, A_1^{00}) \\
\frac{dF^{10}}{dt} = f_F(R^{10}, F^{10}, A_2^{10}), & \frac{dF^{01}}{dt} = f_F(R^{01}, F^{01}, A_2^{01}), & \frac{dF^{00}}{dt} = f_F(R^{00}, F^{00}, A_2^{00}) \\
\frac{dA_1^{10}}{dt} = f_A(R^{10}, F^{10}, A_1^{10}, S_1^{10}), & \frac{dA_1^{01}}{dt} = f_A(R^{01}, F^{01}, A_1^{01}, S_1^{01}), & \frac{dA_1^{00}}{dt} = f_A(R^{00}, F^{00}, A_1^{00}, S_1^{00}), \\
\frac{dA_2^{10}}{dt} = f_A(R^{10}, F^{10}, A_2^{10}, S_2^{10}), & \frac{dA_2^{01}}{dt} = f_A(R^{01}, F^{01}, A_2^{01}, S_2^{01}), & \frac{dA_2^{00}}{dt} = f_A(R^{00}, F^{00}, A_2^{00}, S_2^{00}), \\
\frac{dQ_1^{10}}{dt} = f_Q(A_1^{10}, S_1^{10}, Q(t)), & \frac{dQ_1^{01}}{dt} = f_A(A_1^{01}, S_1^{01}, 0), & \frac{dQ_1^{00}}{dt} = f_A(A_1^{00}, S_1^{00}, 0), \\
\frac{dQ_2^{10}}{dt} = f_Q(A_2^{10}, S_2^{10}, 0), & \frac{dQ_2^{01}}{dt} = f_A(A_2^{01}, S_2^{01}, Q(t)), & \frac{dQ_2^{00}}{dt} = f_A(A_2^{00}, S_2^{00}, 0),
\end{array}
$$

where $Q(t)$ is treated as an exogenous variable whose values are identical to those in the original model. The variant with both pathways removed (right column) is the base case where species $Q$ has no effect on the rotifer. The other two variants (left and middle columns) represent counterfactuals where species $Q$ indirectly affects the rotifer only through one pathway. The equations for Method 1 are similar, with the only difference being that the zeros are replaced with the initial density for species $Q$ for the time interval, i.e., $Q(t_{s-1})$.

### S4 Additional details for the intraguild predation model

#### S4.1 Parameter values and time scaling

All state variables in Hiltunen et al. (2013)'s intraguild predation model are measured in terms of the limiting substrate (N per mL) and time is measured in days. The original and scaled parameter values are listed in Table S1. The scaled parameter values were computed by scaling time by a factor of 1.283 day$^{-1}$. That scaling comes from following the steps in Appendix S1, i.e., it is the maximum ratio of the second and first derivatives for each species ($R_R = 0.888$ day$^{-1}$, $R_F = 1.283$ day$^{-1}$, $R_A = 0.804$ day$^{-1}$, and $R_S = 0.633$ day$^{-1}$). Thus, in the scaled model, unitless time is defined by $\tau = t \times 1.283$.

The effect sizes estimated using the scaled and unscaled models are similar. For the scaled model, Methods 1 and 2, respectively, estimate that the direct effect is 10.5 and 5 times larger on average than the indirect effect. In comparison, for the unscaled model, Methods 1 and 2, respectively, estimate that the direct effect is 8.3 and 3.8 times larger on average than the indirect effect.

Table S1: Original and scaled parameter values for Hiltunen et al. (2013)'s intraguild predation model

| Parameter | Original value | | Scaled value | |
|---|---|---|---|---|
| $g$ | day$^{-1}$ | 0.7 | unitless | 0.546 |
| $k_R$ | N mL$^{-1}$ | 0.25 | N mL$^{-1}$ | 0.25 |
| $\alpha_F$ | unitless | 1 | unitless | 1 |
| $\eta$ | day$^{-1}$ | 1.5 | unitless | 1.169 |
| $\delta$ | day$^{-1}$ | 0.15 | unitless | 0.117 |
| $h$ | N mL$^{-1}$ day$^{-1}$ | 7 | N mL$^{-1}$ | 5.457 |
| $r$ | day$^{-1}$ | 0.7 | unitless | 0.546 |
| $k_A$ | N mL$^{-1}$ | 0.15 | N mL$^{-1}$ | 0.15 |
| $I_f$ | N mL$^{-1}$ day$^{-1}$ | 0.001 | N mL$^{-1}$ | 0.00078 |
| $\alpha_A$ | N mL$^{-1}$ | 2 | N mL$^{-1}$ | 2 |

#### S4.2 Model variants

Method 1 uses four model variants representing all combinations of the two pathways being active or inactive. The following shows the equations for the three model variants where one or both pathways are inactivated; the equations for the variant with two active pathways are identical to model (8) in the main text. For Method 2, the equations for the model variants are identical except that all instances of $R(t_{s-1})$ are replaced with zeros. Note that there is no rotifer equation because rotifer density is treated like an exogenous variable. The blue text highlights the key differences between the model variants.

##### Variant with pathway 1 active and pathway 2 inactive

$$\frac{dF^{10}}{dt} = \frac{hA^{10}F^{10}}{1+\alpha_A A^{10}} - \frac{\eta F^{10} R(t_{s-1})}{k_R + A^{10} + \alpha_F F^{10}} - \delta F^{10} + I_F \tag{S44a}$$

$$\frac{dA^{10}}{dt} = \frac{rS^{10}A^{10}}{k_A + S^{10}} - \frac{gA^{10}R(t)}{k_R + A^{10} + \alpha_F F^{10}} - \frac{hA^{10}F^{10}}{1 + \alpha_A A^{10}} - \delta A^{10} \tag{S44b}$$

$$\frac{dS^{10}}{dt} = \delta - \frac{rS^{10}A^{10}}{k_A + S^{10}} - \delta S^{10} \tag{S44c}$$

**Variant with pathway 1 inactive and pathway 2 active**

$$\frac{dF^{01}}{dt} = \frac{hA^{01}F^{01}}{1 + \alpha_A A^{01}} - \frac{\eta F^{01}R(t)}{k_R + A^{01} + \alpha_F F^{01}} - \delta F^{01} + I_F \tag{S45a}$$

$$\frac{dA^{01}}{dt} = \frac{rS^{01}A^{01}}{k_A + S^{01}} - \frac{gA^{01}R(t_{s-1})}{k_R + A^{01} + \alpha_F F^{01}} - \frac{hA^{01}F^{01}}{1 + \alpha_A A^{01}} - \delta A^{01} \tag{S45b}$$

$$\frac{dS^{01}}{dt} = \delta - \frac{rS^{01}A^{01}}{k_A + S^{01}} - \delta S^{01} \tag{S45c}$$

**Variant with both pathways inactive**

$$\frac{dF^{00}}{dt} = \frac{hA^{00}F^{00}}{1 + \alpha_A A^{00}} - \frac{\eta F^{00}R(t_{s-1})}{k_R + A^{00} + \alpha_F F^{00}} - \delta F^{00} + I_F \tag{S46a}$$

$$\frac{dA^{00}}{dt} = \frac{rS^{00}A^{00}}{k_A + S^{00}} - \frac{gA^{00}R(t_{s-1})}{k_R + A^{00} + \alpha_F F^{00}} - \frac{hA^{00}F^{00}}{1 + \alpha_A A^{00}} - \delta A^{00} \tag{S46b}$$

$$\frac{dS^{00}}{dt} = \delta - \frac{rS^{00}A^{00}}{k_A + S^{00}} - \delta S^{00} \tag{S46c}$$

#### S4.3 Calculations of approximate effect sizes

Using formulas (5) and (6) from the main text, the approximate effects sizes of the direct and indirect effects are,

$$\text{direct effect:} \qquad \frac{1}{2}\frac{\partial f_A}{\partial R} = \frac{-1}{2}\frac{gA}{k_R + A + \alpha_F F}, \tag{S47}$$

$$\text{indirect effect:} \qquad \frac{1}{3!}\frac{\partial f_F}{\partial R}\frac{\partial f_A}{\partial F} = \frac{-1}{3!}\frac{\eta F}{k_R + A + \alpha_F F}\left(\frac{A}{1 + \alpha_A A} - \frac{gAR\alpha_F}{k_R + A + \alpha_F F}\right). \tag{S48}$$

The ratio of those effects is

$$\frac{\text{indirect effect}}{\text{direct effect}} = \frac{-1}{3!}\frac{-\eta F}{3g}\left(\frac{1}{1 + \alpha_A A} - \underbrace{\frac{gR\alpha_F}{k_R + A + \alpha_F F}}_{Q}\right) \approx \frac{-\eta}{3g}\frac{F}{1 + \alpha_A A}. \tag{S49}$$

The approximation in last part of the equation is that the fraction $Q$ is small relative to $\frac{1}{1+\alpha_A A}$. This is true on average for the time series: $Q$ is 40% smaller on average than the fraction $\frac{1}{1+\alpha_A A}$. The approximation is not always valid because $Q$ can be up to 73% of the size of $\frac{1}{1+\alpha_A A}$. However, this only occurs during periods when flagellate density is very low (approximately 5 times smaller than algal density). During those time periods, the indirect effect is approximately 5 time smaller than the direct effect because the indirect effect is proportional to flagellate density while the direct effect is not. Thus, when the approximation is not valid, the indirect effect is much smaller than the direct effect and thus the error in the approximation is irrelevant.

### S5 Additional details for the host-parasite model

#### S5.1 Parameter values and time scaling

The state variables and original and scaled parameters for Strauss et al. (2015)'s host-parasite model are listed in Table S2. The scaled parameter values were computed by scaling time by a factor of 1.082 day$^{-1}$. That scaling comes from following the steps in Appendix S1, i.e., it is the maximum ratio of the second and first derivatives for each state variable ($R_{N_1} = 0.404$ day$^{-1}$, $R_{I_1} = 1.082$ day$^{-1}$, $R_{N_2} = 0.337$ day$^{-1}$, $R_{I_2} = 0.191$ day$^{-1}$, $R_P = 0.297$ day$^{-1}$, and $R_A = 0.669$ day$^{-1}$). Thus, in the scaled model, unitless time is defined by $\tau = t \times 1.082$.

The effect sizes estimated using the scaled and unscaled models are similar. For the scaled model, the ratios of the average effect sizes of the resource-mediated, uptake-mediated, and spore-release pathways are 1:2.03:0.07 and 1:1.47:0.07 for Method 1 and Method 2, respectively. For the unscaled model, the ratios are 1:1.77:0.07 and 1:1.26:0.07 for Method 1 and Method 2, respectively.

Table S2: Variables and original and scaled parameter values for Strauss et al. (2015)'s host-parasite model

| Variable | units* | | | | | |
|---|---|---|---|---|---|---|
| $A$ | $\mu$g chl-$\alpha$ L$^{-1}$ | | | | | |
| $N_i$ | indv. L$^{-1}$ | | | | | |
| $I_i$ | indv. L$^{-1}$ | | | | | |
| $P$ | spore L$^{-1}$ | | | | | |
| Parameter | Original value | | | Scaled value | | |
| | units | host 1 | host 2 | units | host 1 | host 2 |
| $e_i$ | indv. (chi-$\alpha$)$^{-1}$ | 0.189 | 0.687 | indv. (chi-$\alpha$)$^{-1}$ | 0.189 | 0.687 |
| $u_i$ | L indv.$^{-1}$ day$^{-1}$ | 0.021 | 0.00641 | L indv.$^{-1}$ | 0.0194 | 0.0059 |
| $m_i$ | day$^{-1}$ | 0.05 | 0.05 | unitless | 0.0462 | 0.0462 |
| $\mu_i$ | day$^{-1}$ | 0.034 | 0.036 | unitless | 0.031 | 0.033 |
| $\chi_i$ | spores indv. $^{-1}$ | 195000 | 70800 | spores indv. $^{-1}$ | 195000 | 70800 |
| $h_i$ | $\mu$g chi-$\alpha$ L$^{-1}$ | 13.16 | 13.16 | $\mu$g chi-$\alpha$ L$^{-1}$ | 13.16 | 13.16 |
| $\rho_i$ | spore$^{-1}$ | $3.93 \cdot 10^{-4}$ | $3.95 \cdot 10^{-5}$ | spore$^{-1}$ | $3.93 \cdot 10^{-4}$ | $3.95 \cdot 10^{-5}$ |
| $r$ | day$^{-1}$ | 0.9 | | unitless | 0.832 | |
| $K$ | $\mu$g chl-$\alpha$ L$^{-1}$ | 250 | | $\mu$g chl-$\alpha$ L$^{-1}$ | 250 | |
| $\delta$ | day$^{-1}$ | 0.2 | | unitless | 0.184 | |

* indv. = individual; chl-$\alpha$ = chlorophyll-a

#### S5.2 Model variants

Method 1 uses eight model variants representing all combinations of the resource-mediated, uptake-mediated, and spore-release-mediated pathways being active or inactive. The following shows the equations for the model variant where all three pathways are inactivated.

$$\frac{dA^{000}}{dt} = rA^{000}\left(1 - \frac{A^{000}}{K}\right) - u_1 A^{000} N_1^{000} - u_2 A^{000} N_2(t_{s-1}) \tag{S50a}$$

$$\frac{dN_1^{000}}{dt} = e_1 u_1 A^{000} N_1^{000} - m_1 N_1^{000} - \mu_1 I_1^{000} \tag{S50b}$$

$$\frac{dI_1^{000}}{dt} = \rho_1 u_1 [N_1^{000} - I_1^{000}] P^{000} - (m + \mu_1) I_1^{000} \tag{S50c}$$

$$\frac{dI_2^{000}}{dt} = \rho_2 u_2 [N_2(t_{s-1}) - I_2^{000}] P^{000} - (m + \mu_2) I_2^{000} \tag{S50d}$$

$$\frac{dP^{000}}{dt} = \sum_i \chi_i \frac{A^{000}}{h_i + A^{000}} (m_i + \mu_i) I_i^{000} - u_1 P^{000} N_1^{000} - u_2 P^{000} N_2(t_{s-1}) - \delta P^{000} \tag{S50e}$$

Note that there is no equation for host 2 total density because that density is treated like an exogenous variable. The equations for the other variants are identical except that some instances of $N_2(t_{s-1})$ (highlighted in blue) are changed to $N_2(t)$. For example, to get the variant where only the resource-mediated pathways are active, $N_2(t_{s-1})$ in the algal equation is changed to $N_2(t)$ and the superscripts are changed from 000 to 100. Similarly, to get the variant where the resource-mediated and spore-release-mediated pathways are active, $N_2(t_{s-1})$ in the algal equation is changed to $N_2(t)$, $N_2(t_{s-1})$ in the spore equation is changed to $N_2(t)$, and the superscripts are changed from 000 to 101.

For Method 2, the system of equations for each model variant is identical except that all instances of $N_2(t_{s-1})$ are replaced with 0.

#### S5.3 Calculations of approximate effect sizes

Using formulas (5) and (6) from the main text, the approximate effects sizes of the pathways are,

uptake-mediated pathway:

$$\frac{1}{3!}\frac{\partial f_P}{\partial N_2}\frac{\partial f_{I_1}}{\partial P} = \frac{-1}{3!} u_2 P \cdot \rho_1 u_1 S_1, \tag{S51}$$

spore-release-mediated pathway:

$$\frac{1}{4!}\frac{\partial f_{I_2}}{\partial N_2}\frac{\partial f_P}{\partial I_2}\frac{\partial f_{I_1}}{\partial P} = \frac{1}{4!}\rho_2 u_2 P \cdot \frac{\chi_2 A}{h_2 + A}(m_2 + \mu_2) \cdot \rho_1 u_1 S_1 \tag{S52}$$

resource-mediated pathways:

$$\begin{aligned} &\frac{1}{4!}\frac{\partial f_A}{\partial N_2}\frac{\partial f_{N_1}}{\partial A}\frac{\partial f_{I_1}}{\partial A} + \frac{1}{4!}\frac{\partial f_A}{\partial N_2}\frac{\partial f_P}{\partial A}\frac{\partial f_{I_1}}{\partial P} + \frac{1}{5!}\frac{\partial f_A}{\partial N_2}\frac{\partial f_{N_1}}{\partial A}\frac{\partial f_P}{\partial N_1}\frac{\partial f_{I_1}}{\partial P} \\ &= \frac{-1}{4!} u_2 A \Bigg[ e_1 u_1 N_1 \cdot \rho_1 u_1 P + \sum_i \chi_i [m_i + \mu_i] \frac{h_i}{(h_i + A)^2} \cdot \rho_1 u_1 S_1 \\ &\qquad - \frac{1}{5} e_1 u_1 N_1 \cdot u_1 P \cdot \rho_1 u_1 S_1 \Bigg] \qquad \text{(S53)} \\ &\approx \frac{-1}{4!} u_2 A \cdot e_1 u_1 N_1 \cdot \rho_1 u_1 N_1 \qquad \text{(S54)} \end{aligned}$$

The approximation in the formula for the resource-mediated-pathways assumes that the magnitude of the first resource-mediated pathway in Figure 1g is much larger than the magnitudes of the other two resource-mediated pathways; mathematically, the assumption is that $e_1 u_1 N_1 \cdot \rho_1 u_1 P$ is larger than all other terms in the brackets. This assumption is valid at all time points except for a short period around time 25 where the second resource-mediated pathway is larger than the first resource-mediated pathway (details in Appendix S5.4 and Figure S1). However, during that time period, the uptake-mediated pathway is much larger than the both resource-mediated pathways because spore density is much greater than algal density. Thus, while the exact form of the approximation is incorrect for the period around time 25, the rough magnitude of the predicted value is correct because the small algal density is the dominating factor determining the magnitudes of the two resource-mediated pathways. Taking the ratio of the approximate effects for the uptake-mediated and resource-mediated pathways yields equation (11).

#### S5.4 Independent effects of resource-mediated pathways

To estimate the independent effects of the resource-mediated pathways, I follow the steps in Appendix SS.2 and analyze an extended version of the host-parasite model.

Matching the order in Figure 1g, the three resource-mediated pathways are numbered as follows:

$$\begin{aligned}
&\text{Resource-mediated pathway 1:} \quad N_2 \dashv A \to N_1 \to I_1,\\
&\text{Resource-mediated pathway 2:} \quad N_2 \dashv A \to P \to I_1,\\
&\text{Resource-mediated pathway 3:} \quad N_2 \dashv A \to N_1 \to P \to I_1.
\end{aligned}$$

All pathways start with the shared link $N_2 \to A$ and Pathways 1 and 3 also have the shared link $A \to N_1$. Because there are three pathways with overlapping links, I create three copies of host 1 total density ($N_{1,1}$, $N_{1,2}$, $N_{1,3}$) and three copies of the algal density ($A_1$, $A_2$, $A_3$). With these additional variables, the individual pathways are defined as,

$$\begin{aligned}
&\text{Resource-mediated pathway 1:} \quad N_2 \dashv A_1 \to N_{1,1} \to I_1,\\
&\text{Resource-mediated pathway 2:} \quad N_2 \dashv A_2 \to P \to I_1,\\
&\text{Resource-mediated pathway 3:} \quad N_2 \dashv A_3 \to N_{1,3} \to P \to I_1.
\end{aligned}$$

The extended system has three copies of the algal equation and three copies of the equation for host 1 total density. The equations for the extended system are,

$$\frac{dA_i}{dt} = rA_i\left(1 - \frac{A_i}{K}\right) - u_1 A_i N_{1,i} - u_2 A_i N_2, \qquad (i = 1,2,3) \tag{S55a}$$

$$\frac{dN_{1,i}}{dt} = e_1 u_1 A_i N_{1,i} - m_1 N_{1,i} - \mu_1 I_1, \qquad (i = 1,2,3) \tag{S55b}$$

$$\frac{dI_1}{dt} = \rho_1 u_1 [N_{1,1} - I_1] P - (m + \mu_1) I_1, \tag{S55c}$$

$$\frac{dI_2}{dt} = \rho_2 u_2 [N_2 - I_2] P - (m + \mu_2) I_2, \tag{S55d}$$

$$\frac{dP}{dt} = \sum_i \chi_i \frac{A_2}{h_i + A_2}(m_i + \mu_i) I_i - u_1 P N_{1,3} - u_2 P N_2 - \delta P. \tag{S55e}$$

I note the following about the extended system. First, there is no equation for host 2 total density ($dN_2/dt$), because $N_2$ is treated like an exogenous variable in all model variants. Second, the equations for each algal density ($A_i$) and host 1 total density ($N_{1,i}$) only depend on their respective variables, e.g., the $dA_1/dt$ and $dN_{1,1}/dt$ equations only depend on $A_1$ and $N_1$ and do not depend on $A_2$, $A_3$, $N_2$, or $N_3$. This ensures that the three pathways are separated. Third, all other equations depend on the variables associated with their specific pathways. In particular, the equation for host 1 infected density ($dI_1/dt$) depends on $N_{1,1}$ because only pathway 1 includes a direct effect on host 1 total density on host 1 infected density. The equation for spore density ($dP/dt$) depends on both $A_2$ and $N_{1,3}$. The spore release terms depend on $A_2$ because pathway 2 contains the direct effect of algal density on spore density; the uptake term depends on $N_{1,3}$ because pathway 3 contains the direct effect of host 1 total density on spore density. Combined, these properties allow for the individual effects of each pathway to be estimated.

Estimating the individual effects of the uptake-mediated pathway, spore-release-mediated pathway, and three resource-mediated pathways requires $2^5 = 32$ model variants. The variables of the variants are denoted using superscripts $p_1p_2p_3p_4p_5$ where $p_1$, $p_2$, and $p_3$ are the indicators for the three resource-mediated pathways, $p_4$ is the indicator for the uptake-mediated pathway, and $p_5$ is the indicator for the spore-release-mediated pathway. For example, $A^{10010}$ is a variable in the model variant where the first resource-mediated pathway and the uptake-mediated pathway are present and the other three pathway are absent. For Method 1, the model variant with all inactive pathways has equations,

$$\frac{dA_1^{00000}}{dt} = rA_1^{00000}\left(1 - \frac{A_1^{00000}}{K}\right) - u_1A_1^{00000}N_{1,1}^{00000} - u_2A_1^{00000}N_2(t_{s-1}), \tag{S56a}$$

$$\frac{dA_2^{00000}}{dt} = rA_2^{00000}\left(1 - \frac{A_2^{00000}}{K}\right) - u_1A_2^{00000}N_{1,2}^{00000} - u_2A_2^{00000}N_2(t_{s-1}), \tag{S56b}$$

$$\frac{dA_3^{00000}}{dt} = rA_3^{00000}\left(1 - \frac{A_3^{00000}}{K}\right) - u_1A_3^{00000}N_{1,3}^{00000} - u_2A_3^{00000}N_2(t_{s-1}), \tag{S56c}$$

$$\frac{dN_{1,1}^{00000}}{dt} = e_1u_1A_1^{00000}N_{1,1}^{00000} - m_1N_{1,1}^{00000} - \mu_1I_1^{00000}, \tag{S56d}$$

$$\frac{dN_{1,2}^{00000}}{dt} = e_1u_1A_2^{00000}N_{1,2}^{00000} - m_1N_{1,2}^{00000} - \mu_1I_1^{00000}, \tag{S56e}$$

$$\frac{dN_{1,3}^{00000}}{dt} = e_1u_1A_3^{00000}N_{1,3}^{00000} - m_1N_{1,3}^{00000} - \mu_1I_1^{00000}, \tag{S56f}$$

$$\frac{dI_1^{00000}}{dt} = \rho_1u_1[N_{1,1}^{00000} - I_1^{00000}]P^{00000} - (m+\mu_1)I_1^{00000}, \tag{S56g}$$

$$\frac{dI_2^{00000}}{dt} = \rho_2u_2[N_2(t_{s-1}) - I_2^{00000}]P^{00000} - (m+\mu_2)I_2^{00000}, \tag{S56h}$$

$$\frac{dP^{00000}}{dt} = \sum_i \chi_i\frac{A_2^{00000}}{h_i + A_2^{00000}}(m_i+\mu_i)I_i^{00000} - u_1P^{00000}N_{1,3}^{00000} - u_2P^{00000}N_2(t_{s-1}) - \delta P^{00000}. \tag{S56i}$$

The equations for the other variants are identical except that specific instances of $N_2(t_{s-1})$

(highlighted in blue) are changed to $N_2(t)$. For Method 2, the system of equations for each model variant is identical except that all instances of $N_2(t_{s-1})$ are replaced with 0.

Figure S1 shows the estimated individual effects for the five pathways. The effects sizes for the spore-release-mediated (red) and uptake-mediated (cyan) pathways are the same as Figure 3. The effect sizes of the three-resource mediated pathways (blue, purple, green) sum up to the magnitude of the aggregated effect (blue) in Figure 3.

For both Methods 1 and 2, resource-mediated pathway 1 is much larger on average than the other two resource-mediated pathways. Specifically, on average, the ratios of the magnitudes for the three resource-mediated pathways are 1:0.25:0.007 for Method 1 and 1:0.25:0.01 for Method 2. However, the effect of resource-mediated pathway 2 is larger than the effect of resource-mediated pathway 1 for a short period around time 25 (compare blue and magenta curves). This reversal can be explained by computing the approximate effect sizes using equation (6),

$$\frac{\text{Pathway 1}}{\text{Pathway 2}} = \frac{\frac{1}{5!}\frac{\partial f_A}{\partial N_2}\frac{\partial f_{N_1}}{\partial A}\frac{\partial f_{I_1}}{\partial N_1}}{\frac{1}{5!}\frac{\partial f_A}{\partial N_2}\frac{\partial f_P}{\partial A}\frac{\partial f_{I_1}}{\partial P}} = \frac{e_1 u_1 N_1 \cdot \rho_1 u_1 P}{[\chi_1(m_1+\mu_1)I_1 + \chi_2(m_2+\mu_2)I_2]\frac{h_1}{[h_1+A]^2} \cdot \rho_1 u_1 [N_1 - I_1]} \tag{S57}$$

$$= \frac{N_1}{N_1 - I_1}\frac{(h_1+A)^2}{h_1}\frac{e_1 u_1 N_1 P}{\chi_1(m_1+\mu_1)I_1 + \chi_2(m_2+\mu_2)I_2}. \tag{S58}$$

Near time 25, most hosts are susceptible ($N_1 - I_1 \approx N_1$), algal density is lower, and spore densities are lower. Consequently, the first fraction is close to 1, the second fraction is close to 25, and the third fraction is very small because host 1's maximum burst size ($\chi_1$) is very large compared to the host total 1 density and spore density. As a result, resource-mediated pathway 2 has a larger magnitude than resource-mediated pathway near time 25. However, later in time, most hosts are infected ($N_1 - I_1$ much smaller than $N_1$) and algal density is high, and spore density is much larger than the maximum burst size for host 1. Consequently, all fractions are larger than 1 in magnitude and resource-mediated pathway 1 has a larger magnitude than resource-mediated pathway 2. The biological interpretation of this is the following. Near time 25, algal densities are low and resource competition primarily reduces host 1 disease levels by limiting replication of the pathogen within infected hosts, leading to reduced burst sizes. Later time time when algal densities are higher, resource competition primarily reduces host 1 disease levels by limiting host 1's reproduction, which limits the number of susceptible individuals that can become infected.

When comparing all five pathways, the uptake-mediated pathway has the largest effect, resource-mediated pathway 1 has the next largest, and the other pathways have much smaller effects. In particular, the ratios of the average effect sizes of the three resource-mediated, uptake-mediated, and spore-release pathways are 1:0.25:0.007:2.53:0.09 for Method 1 and 1:0.25:0.01:1.76:0.09 for Method 2. Note that while initially smaller, the effect of resource-mediated pathway 1 is comparable in magnitude to the effect of the uptake-mediated pathway near the end of the time series; this was also observed in the analysis with the aggregated effects of competition.

The 2-way, 3-way, 4-way, and 5-way interactive effects are not show in Figure S1 because they are at least an order of magnitude smaller than the individual effects. The one exception is the 2-way interactive effect between resource-mediated pathway 1 and the uptake-mediated pathway (Figure S1e). That two-way interactive effect is larger in magnitude than the other three pathways (resource-mediated pathways 1 and 2 and spore-release-mediated pathway). However, because it is less than five times smaller than the individual effects of the resource-mediated 1 and uptake-mediated pathways, it does not qualitatively change the biological interpretations of the results.

Combining these results and those of the main text, both methods predict that spore uptake by host 2 (cyan curve in Figure S1) plays the largest role in decreasing host 1 infected density. However, late in time, spore uptake and resource consumption have effects of comparable magnitude. During that time, the effects of resource competition are primarily driven by reductions in host reproduction (blue curve in Figure S1). The effects of the other resource-mediated pathways and spore-release-mediated pathway are minor because they are always small in magnitude compared to the effect of the uptake-mediated pathway.

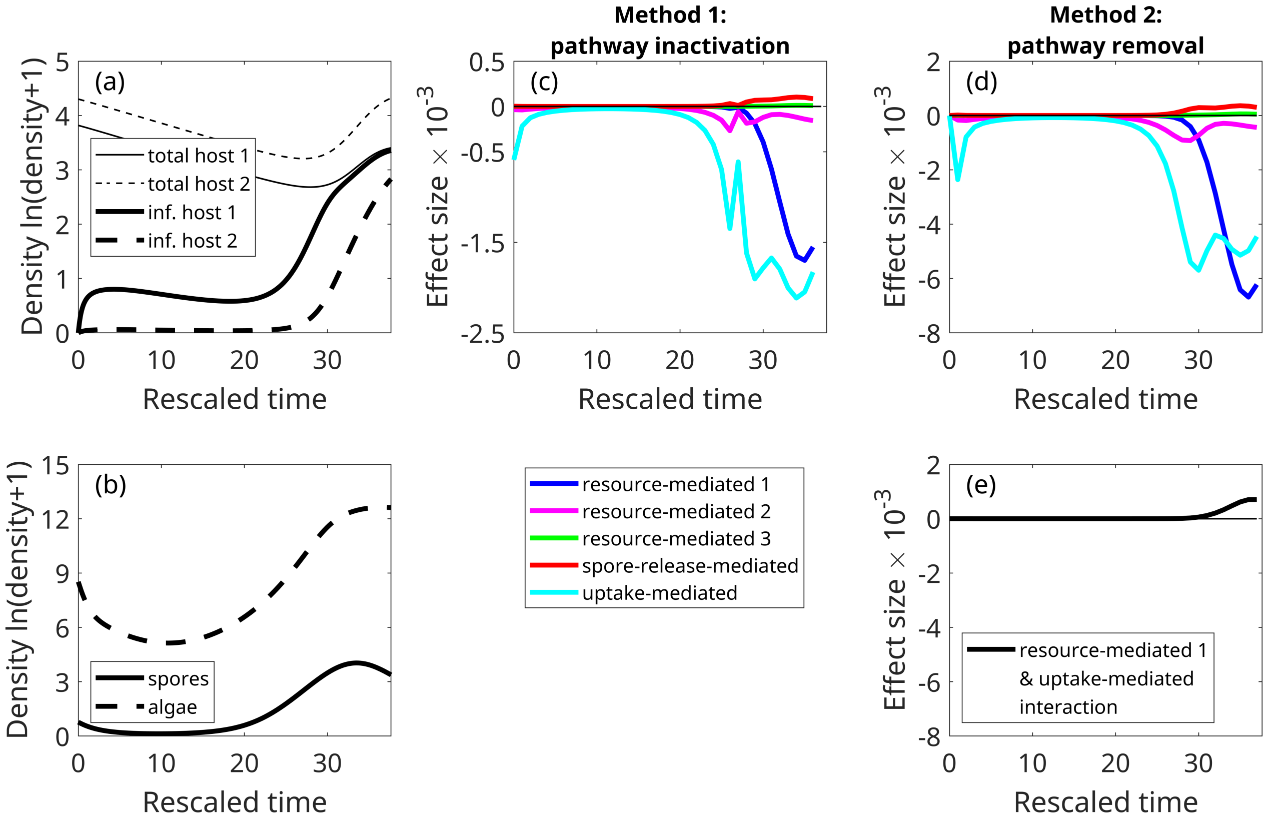

Figure S1: Estimated effect sizes for the three resource-mediated, spore-release-mediated, and uptake-mediated pathways. (a) Time series of total densities (thin curves) and infected densities (thick curves) for host 1 (solid curves) and host 2 (dashed curves) generated by model (1). (b) Accompanying time series of spore density (solid) and algal density (dashed). Time series of individual effects estimated by (c) Method 1 and (d) Method 2. The numbering of the resource-mediated pathways matches the order in Figure 1g; see Appendix S5.3 for more details. (e) Two-way interactive effect of the resource-mediated 1 pathway and the uptake pathway; all other interactive effects are not shown because they are at least an order of magnitude smaller than the independent effects.
